# Targeted epigenetic repression of oncogenic transcription factors via CRISPR/dCas9 locus-specific silencing

**DOI:** 10.64898/2026.06.27.734664

**Authors:** Shahama Taifour, Christopher Wallis, Edina Wang, Eleanor Woodward, Ash W.J. Tie, Charlene Waryah, Larissa Dymond, Andrew Woo, Peter Houghton, K Swaminathan Iyer, Marck Norret, Cameron W Evans, Louise Winteringham, Silvana Gaudieri, Pilar Blancafort

## Abstract

Despite the revolutionary impact of genome engineering tools in medicine, the safe and effective intracellular delivery of CRISPR remains a major obstacle for clinical applications. Here, we implement precision molecular medicine and delivery strategies based on CRISPR/dCas9 systems adapted for epigenetic repression (dCas9-KRAB) to silence oncogenic drivers with high genomic selectivity. As proof-of-principle, we target the *EWSR1-FLI1* translocation, which encodes a chimeric and hard-to-drug oncogenic transcription factor driving approximately 85% of the cases of Ewing Sarcoma (EWS)-an aggressive malignancy affecting children and adolescents. We describe the development of a non-viral and programmable polymeric system for the delivery of dCas9-KRAB as ribonucleoprotein (RNP) payloads for selective *EWSR1-FLI1* repression. We demonstrate highly efficient intracellular delivery of RNPs loaded in polyamide-amine (PAMAM) polymers functionalized by guanidino groups, resulting in robust silencing of *EWSR1-FLI1* both in established cell line xenografts and in patient-derived xenografts (PDXs) of EWS. Moreover, silencing of *EWSR1–FLI1* is accompanied by potent anti-tumor effects. To our knowledge, we describe the first non-viral platform for *in vivo* delivery of dCas9-KRAB/RNPs, which can be adapted for the repression of any oncogene. We further outline dCas9/RNP formulations for future therapeutic applications to treat poor-prognosis cancers driven by hard-to-drug oncogenes.

## Introduction

The clustered Regularly Interspaced Short Palindromic Repeats/Cas9 nuclease (CRISPR/Cas9) system of *Streptococcus pyogenes* is a powerful, versatile, and highly efficient tool for genome engineering.^1–3^ In this system, endonuclease enzyme Cas9 specifically interacts with single guide RNA (gRNA), which directs Cas9 to the target genomic sequence.^4–7^ CRISPR technologies have been rapidly expanded by repurposing the Cas9 enzyme to manipulate the cancer epigenome,^8–11^ providing alternative methods for highly effective gene silencing without causing irreversible changes in the DNA, and with the possibility of maintaining repression of all different isoforms of a given oncogene driven by the same promoter.^12^

In the CRISPR-based epigenome editing systems, ‘nuclease-dead’ Cas9 (dCas9) is a catalytically deactivated version of Cas9 endonuclease in which both catalytic domains (HNH and RuvC) are mutated by site-directed mutagenesis. Although these mutations maintain the DNA-binding specificity of Cas9, they abolish wild-type Cas9 endonuclease activity.^13–15^ The resulting dCas9 protein can be tethered to different modular effector domains in order to alter the epigenetic state of targeted genes, typically in regulatory regions such as enhancers and promoters, resulting in transcriptional perturbations.^13,15,16^ Notably, linking dCas9 to a potent transcriptional repressor domain, the Krüppel-associated box (KRAB), creates a compact and effective dCas9 artificial repressor system, CRISPR/dCas9-KRAB, which has demonstrated negligible off-target activities.^11,13,17–19^ The transcriptional silencing by the KRAB domain involves the recruitment of the KRAB-associated protein 1 (KAP1) co-repressor, which associates with a “cloud” of epigenetic silencers, including the H3K9 methyltransferase SETDB1, the heterochromatin protein 1 alpha (HP1α) and the histone deacetylase NURD complex.^20–23^ When dCas9-KRAB is directed to the regulatory regions of targeted genes, it promotes a reduction in H3K27 and H3K9 acetylation, and, reciprocally, an increase in H3K9 and H3K27 tri-methylation, resulting in heterochromatinization and transcriptional repression of target genes (Figure 1A).^13,24,25^

**Figure 1.**
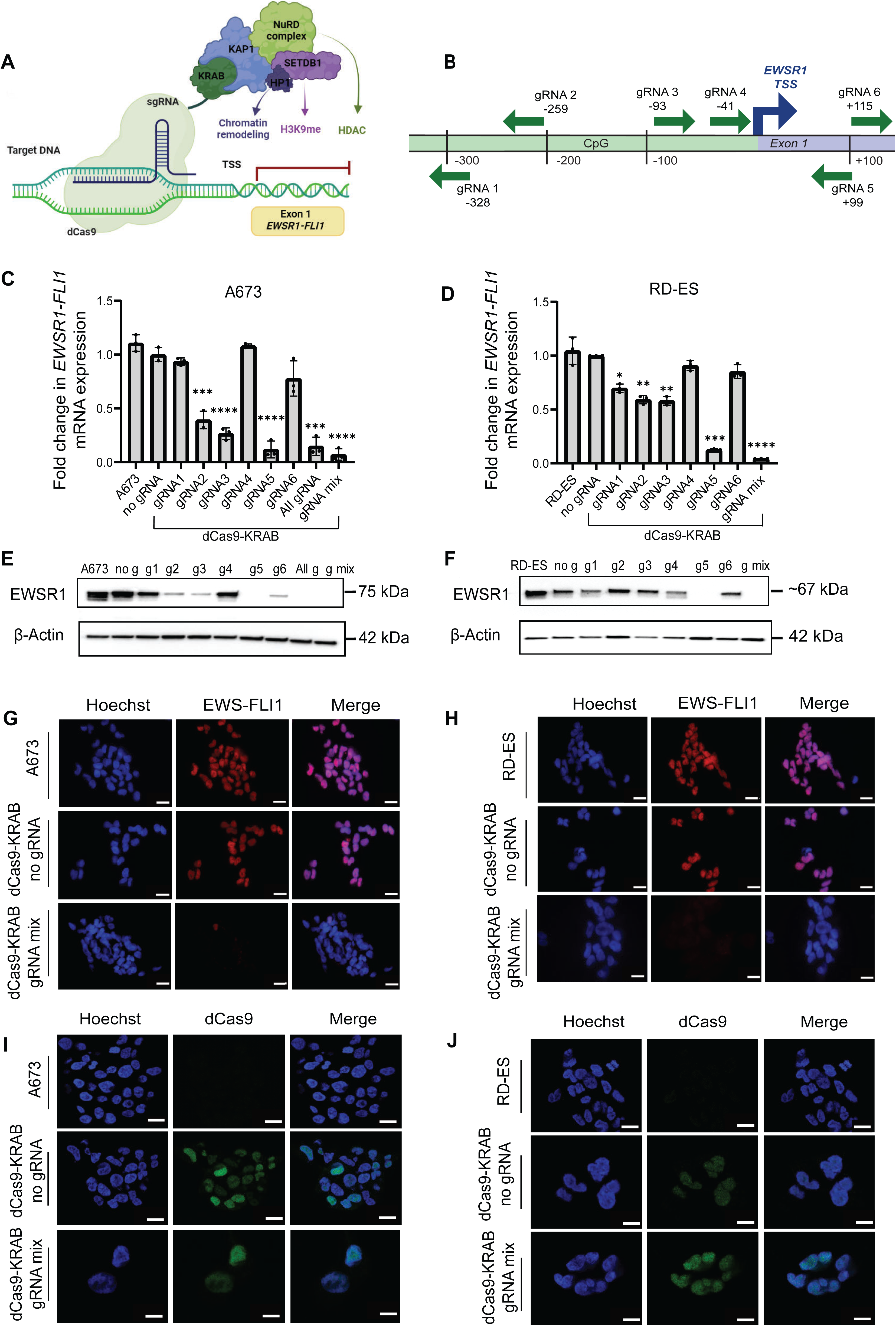
Repression of *EWSR1-FLI1* oncogene using dCas9-KRAB in A673 and RD-ES cells. (A) Schematic representation of the dCas9-KRAB system. dCas9-KRAB is directed to the target promoter via different gRNAs tiled in the promoter and first exon of *EWSR1*. KRAB interacts with the KRAB domain-associated protein 1 (KAP-1) co-repressor, which then recruits histone-modifying enzymes to induce heterochromatin formation: H3K9 methyltransferase SET Domain Bifurcated 1 (SETDB1); heterochromatin protein 1 (HP1); and nucleosome remodeling deacetylase (NuRD) complex. This leads to the loss of H3K27ac and H3K9ac, while increasing H3K9me3 and H3K27me3, ultimately stimulating transcriptional repression. (B) The positions of the designed six gRNAs targeting the *EWSR1* proximal promoter and the associated 5′ untranslated regions. Green arrows show gRNAs pointing 5’ to 3’ orientation. The position of each gRNA is numbered according to the distance from the transcription start site of the *EWSR-FLI1* mRNA transcript variant 2 (NM_005243.4). (C,D) qRT-PCR analysis of *EWSR1-FLI1* mRNA expression in A673 (C) and RD-ES (D) cells normalized to dCas9-KRAB with no gRNA sample, with *GAPDH* as a housekeeping gene. Data are presented as mean ± standard deviation. *n* = 3 biologically independent experiments. *p* values were determined by the Student’s *t*-test compared to dCas9-KRAB no gRNA samples (\**p* < 0.05, \*\**p* < 0.01, \*\*\**p* < 0.001, \*\*\*\**p* < 0.0001). *p* values were determined by the Student’s *t*-test for gRNA mix compared to gRNA 5 (\*\**p* < 0.01). Comparisons not labeled were not statistically significant (*p* > 0.05). (E,F) Representative western blots for the detection of EWS-FLI1 protein and β-Actin in A673 (E) and RD-ES (F) cell lines. (G,H) Immunostaining of EWS-FLI1 (Red) and Hoechst staining (blue) in A673 (G) and RD-ES (H) cell lines. Scale bars indicate 50 μm. I,J) Immunostaining of dCas9 (green) and Hoechst staining (blue) in A673 (I) and RD-ES (J) cell lines. Scale bars indicate 10 μm.

While CRISPR technology offers avenues for precision medicine,^11^ safe and effective delivery remains a hurdle for successful clinical translation.^26–28^ Delivery systems based on viral vehicles are limited by their packaging capacity, immunogenicity, and potential risk of insertional mutagenesis.^29^ Non-viral vectors have been developed to overcome these limitations; however, they are still intrinsically deficient in combining highly efficient delivery with low cytotoxicity.^30^ The delivery of Cas9 and gRNA into targeted cells as non-viral formulations, either as plasmid DNA or mRNA/gRNAs, has shown limitations due to unwanted off-target effects or extremely low stability.^31–33^

The dCas9/gRNA ribonucleoprotein (dCas9-KRAB RNP) complex is an attractive approach due to its low immunogenicity, high on-target specificity, negligible off-target effects, and absolute clearance of dCas9 post-transduction of the cells/tissues.^33–35^ Nevertheless, targeting tumor tissues with dCas9-KRAB RNPs remains a major challenge. In this context, the large size of dCas9 fused to epigenetic effectors, the endosomal entrapment of dCas9 and gRNA, and the degradation/denaturation of these large dCas9 molecules inside the cells represent substantial barriers to *in vivo* delivery.^36,37^ Furthermore, producing stable nanocarriers that effectively deliver these RNPs to targeted organs within the body still remains a significant problem for the CRISPR field.^38^

The architecture of non-viral agents can be dynamically manipulated or functionalized to achieve efficient cytosolic delivery of both protein and RNA payloads.^37^ To this aim, diverse synthetic vehicles, such as lipid nanoparticles, inorganic nanoparticles, and cationic polymers, have been developed.^39^ While recent works outlined lipid and polymeric nanocarriers for *in vivo* delivery of Cas9 RNPs, to the best of our knowledge, no report has shown successful delivery of dCas9-KRAB silencers in solid tumors and patient derived xenografts^40–43^.

Recently, we described highly effective dendronized polymers for the delivery of nucleic acids *in vitro* and *in vivo*.^44^ This polymeric formulation combines high flexibility with lower charge density to reduce cytotoxicity and enhance cargo packaging density.^44^ Later, we reported that partial functionalization of 5G dendronized polymer surface with *p*-guanidinobenzoic acid (hereafter referred to as GBA-polymer) offered exceptional protein delivery by protecting the payload from endosomal degradation and permitting controlled endosomal release of proteins.^37^

Herein, we assessed the therapeutic efficacy of these polymeric formulations for dCas9-KRAB RNP delivery in a solid tumor model of Ewing sarcoma (EWS), a highly aggressive bone and soft tissue malignancy occurring in children and young adults. While accounting for 2% of all pediatric cancers^45,46^, these cancers are often resistant to chemotherapy, disease recurrence and metastases, reflected in a 5-year overall survival rate of less than 30%.^47,48^ Nearly 85% of EWS cases harbor the t(11;22)(q24;q12) chromosomal translocation that produces the *EWSR1-FLI1* oncogenic fusion.^49^ The resulting *EWSR1-FLI1* chimera encodes a neomorphic transcription factor (TF) with enhanced transcriptional activity in neoplastic cells. Like many other TFs, EWS-FLI1 has intrinsically disordered structures that lack small molecule binding pockets and enzymatic activity, rendering them refractory to effective drug design.^50,51^ The EWSR1-FLI1 TF reprograms the transcriptional and epigenetic landscape of EWS either by generating *de novo* enhancers at GGAA repeat elements or by displacing ETS transcription factors at ETS-like sequences (single GGAA motif).^52,53^ As a result, the EWS-FLI1 oncoprotein functions as an autonomous, cancer specific switch, whose persistent expression is essential for tumorigenesis, representing an excellent model system to assess the impact of EWS-FLI1 silencing *via* dCas9-KRAB RNPs. ^54^

This manuscript reports the development of a non-viral polymeric delivery system for CRISPR/RNP payloads and demonstrates therapeutic efficacy both *in vitro* and *in vivo*. In addition, our platform shows promising results in preclinical EWS patient-derived xenograft (PDX) models. Given the versatile nature of the RNP systems, which rely on synthetic short RNAs that can be quickly repurposed to other targets of interest, we describe a platform which can be harnessed for the repression “at will” of any hard-to-drug oncogene in solid tumors.

## Results

### CRISPR/dCas9-KRAB specifically and efficiently suppresses *EWSR1-FLI1* in EWS cell lines using lentiviral delivery systems

To silence the endogenous *EWSR1-FLI1* fusion, we selected two human EWS cell lines (A673 and RD-ES) harboring the *EWSR1-FLI1* chromosomal translocation.^55,56^ We took advantage of the *S. pyogenes* CRISPR system in which dCas9 is C-terminally fused to the KRAB repressor domain (*Sp*dCas9-KRAB, Figure 1A).^24,57^ To identify optimal gRNAs targeting the *EWSR1-FLI1* fusion, we designed six gRNAs encompassing the *EWSR1* proximal promoter and the associated 5′ untranslated regions (UTR) using the Benchling CRISPR design computational tool (Figure 1B; and Table S1, Supporting Information 1).^58–60^

Each gRNA was cloned into the dCas9-KRAB vector and delivered into both EWS cell lines with the pLV lentiviral vector system in which both gRNA and dCas9-KRAB are co-expressed in the same vector.^24^ Real-time quantitative reverse transcriptase PCR (qRT-PCR) was performed to investigate the silencing capacity of single gRNAs, a mix of all six gRNAs, and a specific combination of the most effective gRNAs. The dCas9-KRAB in absence of gRNA (no gRNA sample, Figure 1C) was used as a control for normalization.

In the A673 cell line, the expression of *EWSR1-FLI1* mRNA was significantly suppressed by gRNA 2 (0.39-fold, *p* = 0.0005), gRNA 3 (0.26-fold, *p* = 0.0001), and gRNA 5 (0.12-fold, *p* = 0.0001) relative to dCas9-KRAB with no gRNA. Similarly, in the RD-ES cells, *EWSR1-FLI1* mRNA expression was significantly suppressed by gRNA 2 (0.59-fold, *p* = 0.004), gRNA 3 (0.58-fold, *p* = 0.004), and gRNA 5 (0.1-fold, *p* = 0.0002) relative to dCas9-KRAB with no gRNA.

The combined use of the three most potent gRNAs (gRNAs 2, 3, and 5; hereafter referred to as “gRNA mix”) demonstrated superior suppression of *EWSR1-FLI1* compared to the single gRNAs. Remarkably, gRNA mix achieved the highest repression of *EWSR1-FLI1* compared to gRNA 5, particularly in RD-ES (0.044-fold, *p* = 0.0041). In addition, the repression of *EWSR1-FLI1* by gRNA mix was superior to the repression observed when combining all six gRNAs (0.15-fold, *p* = 0.0002) in A673 cells (Figure 1C,D).

To validate the targeting specificity of dCas9-KRAB-mediated repression using the three best gRNAs, we computationally generated gRNA sequences comprising three or fewer mismatches relative to all the cognate *EWSR1-FLI1* gRNA sequences, which mapped into the proximal promoter of sixteen putative “off-target” genes (Table S2, Supporting Information 1). As observed with other recent reports that used CRISPR/dCas9 systems, no significant off-target gene regulation was observed for the sixteen genes in A673 cells transduced with dCas9-KRAB with gRNA mix relative to dCas9-KRAB with no gRNA, as assessed by qRT-PCR (Figure S1, Supporting Information 1).^61,62^

Next, western blotting (Figure 1E,F; and Figure S2, Supporting Information 1) and immunofluorescence experiments (IF) (Figure 1G,H) were performed to detect EWSR1 protein expression. The immunoblots in WT cells showed a main band of ∼67 and 75 kDa corresponding to the reported molecular weight of the fusion in A673 and RD-ES, respectively, and also a faster migrating band which could correspond to an alternative isoform of the fusion.^63^ These results confirmed the qRT-PCR data and demonstrated the nearly complete silencing of *EWSR-FLI1,* particularly with gRNA mix at the protein level by the gRNA mix in both cell lines, affecting all possible isoforms of the fusion. Furthermore, the expression of dCas9-KRAB in A673 and RD-ES in the transduced cells was verified using IF staining and compared to non-transduced control (Figure 1I,J). Efficient transduction of both cell lines was also confirmed using an EGFP-expressing lentiviral vector (Figure S3, Supporting Information 1). Overall, these data demonstrated that the combination of three gRNAs with dCas9-KRAB was able to achieve highly effective silencing of the endogenous *EWSR1-FLI1* oncogene in representative EWS cell lines.

### Silencing of oncogenic *EWSR1-FLI1* attenuates cell proliferation, migration, and colony formation in EWS cell lines

Since the *EWSR1-FLI1* translocation is the main driving oncogene leading to malignant transformation and metastatic progression in EWS,^64^ we next explored the phenotypic consequences of *EWSR1-FLI1* silencing by dCas9-KRAB after epigenetic editing.

We first monitored changes in cell viability over time in EWS cell lines transduced with dCas9-KRAB silencing constructs and controls. We found that dCas9-KRAB with gRNA mix significantly decreased cell viability after 96 h of cell seeding (*p* < 0.0001) in A673 and RD-ES, respectively, relative to no gRNA (Figure 2A,B). As expected, the silencing of *EWSR1-FLI1* dramatically abolished anchorage-independent cell growth and clonogenicity. The number of colonies in soft agar was significantly reduced in A673 (0.23%, *p* < 0.0001) and RD-ES (29.6%, *p* = 0.01) cell lines when dCas9-KRAB was delivered with gRNA mix compared to no gRNA (Figure 2C,D).

**Figure 2.**
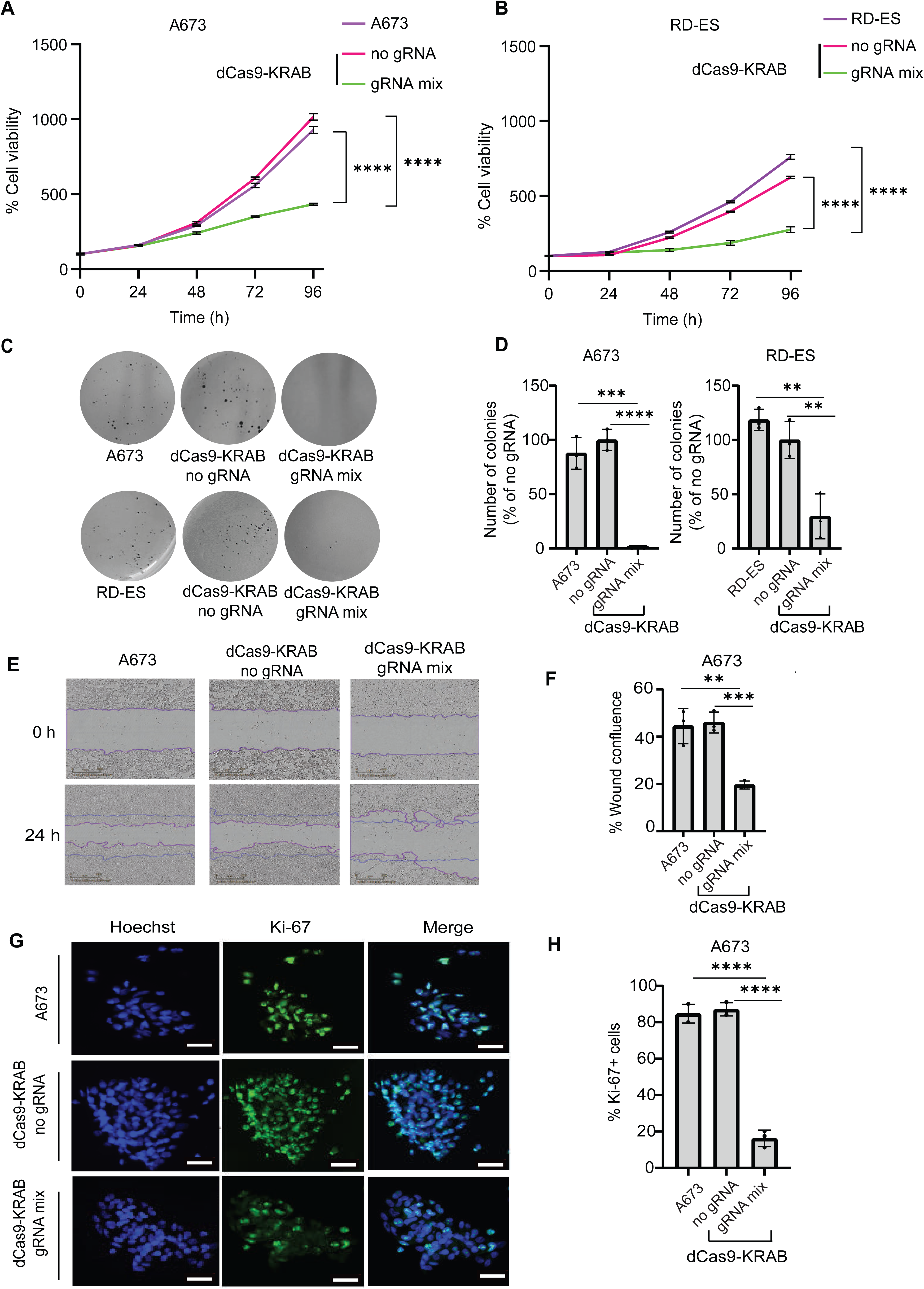
Cellular phenotypic alterations induced by *EWSR1-FLI1* silencing using dCas9-KRAB off-switches in EWS cell lines. (A,B) Decreased cell viability post-*EWSR1-FLI1* silencing in A673 and RD-ES cells, respectively. Cell viability was measured every 24 h for a total of 96 h using the CellTiter-Glo® 2.0 Assay. Data are shown as mean values ± SD of three independent experiments; \*\*\*\**p* < 0.0001. (C) Representative images of a clonogenic assay 2-3 weeks after seeding A673 and RD-ES cells in soft agar. (D) Quantification of soft agar colony formation. Data are normalized to dCas9-KRAB with no gRNA and displayed as mean values ± SD of three independent experiments; \*\*\*\**p* < 0.0001; \*\*\**p* < 0.001 and \*\**p* < 0.01. (E) Representative images of migrating A673 cells in the scratched area using the IncuCyte ZOOM system from Essen Bioscience (magnification 10×). (F) Relative wound confluence is shown at 24 h post-seeding of the cells. \*\**p* < 0.01, \*\*\**p* < 0.001; n = 3; error bars represent SD. (G) Representative immunofluorescence staining of Ki-67 (green) and nuclei (Hoechst 33258, blue). (H) Quantification of Ki-67 positive cells by immunofluorescence. Scale bar represents 100 μm. Data are presented as mean values ± SD of three independent experiments; \*\*\*\**p* < 0.0001.

Next, the effect of *EWSR1-FLI1* silencing on cell migration was investigated in a scratch wound-healing assay. Epigenetic editing of *EWSR1-FLI1* was associated with decreased wound confluence after 24 h when dCas9-KRAB was delivered with gRNAs (19.5%, *p* < 0.001) compared to no gRNA control (43%) (Figure 2E,F). As expected, together with changes in migratory behavior, the *EWSR1-FLI1* silencing by dCas9-KRAB with gRNA mix also induced changes in cell morphology and reorganization of the actin cytoskeleton, resulting in longer and thicker F-actin stress fibers with larger-spread shape compared to that for no gRNA and control (Figure S4, Supporting Information 1). Our reported changes in migration are consistent with another study using RNAi methods to target the *EWSR1-FLI1* oncogenic fusion in TC71 cells.^65^

Finally, we conducted IF staining for the Ki-67 cell proliferation antigen to quantify changes in tumor cell proliferation after epigenetic editing. We detected a significantly lower frequency of actively proliferating cells (Ki-67-positive cells) after cellular transduction with dCas9-KRAB with gRNA mix (14.4%, *p* < 0.0001) compared to that for no gRNA (88.6%) in A673 cells (Figure 2G,H). In contrast to EWS cells, HDFa cells (normal primary human fibroblasts not encoding the oncogenic fusion) transduced with the same CRISPR repressive constructs to downregulate the *EWSR1* promoter did not show significant changes in cell proliferation when comparing CRISPR-edited cells vs. controls, as assessed by Ki-67 staining. This indicated that EWS tumor cells harboring the *EWSR1-FLI1* fusion were dependent on the expression of the oncogenic TF to maintain cell proliferation, whereas non-transformed cells not carrying the fusion were not as reliant on WT *EWSR1* activity to support proliferation (Figures S5 and S6, Supporting Information 1). Our data is consistent with a recent RNAi study which demonstrates that downregulation of *EWSR1* in non-EWS cell lines does not significantly perturb cell proliferation.^66^

Taken together, these results demonstrated that suppression of *EWSR1-FLI1* expression in EWS cells by dCas9-KRAB resulted in decreased cell survival and proliferation, and impaired migratory capacity and tumorigenicity. Importantly, our data also suggest that downregulating *EWSR1* in “normal” diploid cells would not cause significant toxicity to healthy cells.

### CRISPR/dCas9-KRAB-mediated silencing of *EWSR1-FLI1* induces transcriptome reprogramming in EWS cells

To investigate the genome-wide transcriptomic changes induced by dCas9-KRAB, we performed bulk RNA-sequencing (RNA-seq) by comparing A673 cells transduced with dCas9-KRAB with the active mix of gRNAs with dCas9-KRAB no gRNA or with non-transduced cells, using biological triplicates. Non-transduced cells were profiled to subtract background due to lentiviral transduction (Figure 3).

**Figure 3.**
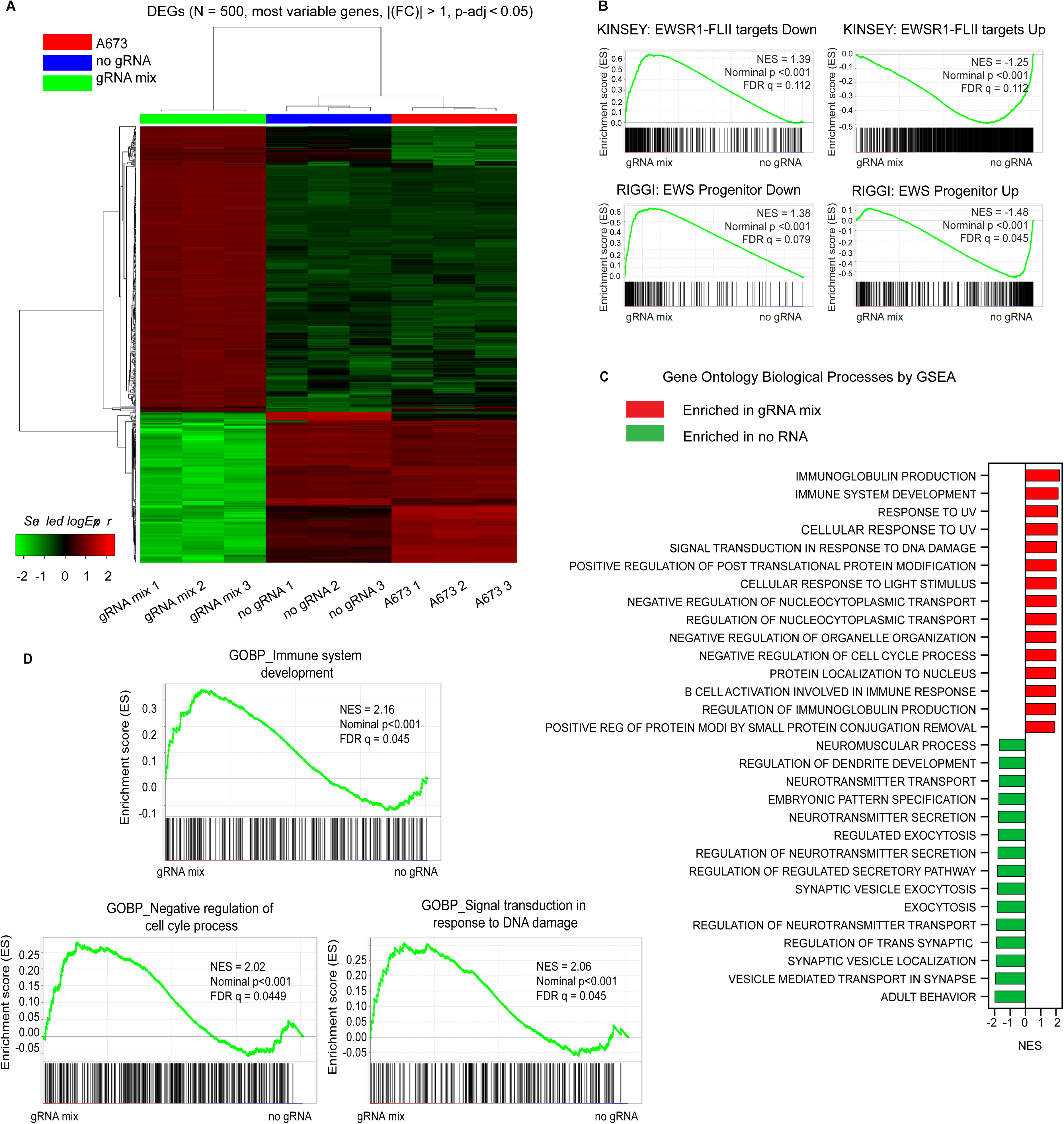
Bioinformatic analysis of differentially expressed genes based on RNA sequencing data. (A) Hierarchical clustering heatmap of the top 500 differentially expressed genes (DEGs) for A673 cells transduced with dCas9-KRAB with gRNA mix compared to no gRNA cells and A673 control, using iDEP.96 software. The horizontal axis displays samples name, while the vertical axis depicts the Fold Change (FC) of the top 500 most significant DEGs. The upregulated genes are in red, and the downregulated genes are in green. Sample types are shown in the color bar above: blue for A673 control cells, green for no gRNA samples, and red for gRNA mix cells. (B) Gene set enrichment analysis (GSEA) plot of *EWSR1-FLI1* target genes, reported by Kinsey *et al.* in TC71 and EWS502 cells post *EWSR1-FLI1* knockdown by shRNA, in A673 cells by dCas9-KRAB with gRNA mix compared with no gRNA samples (top). GSEA plots of up- and downregulated genes, induced by ectopic expression of EWS-FLI1 in human MSCs, in A673 cells by dCas9-KRAB with gRNA mix compared with no gRNA samples (bottom). The y-axis represents the enrichment score (ES) for individual genes. The x-axis represents the gene set enrichment. NES, normalized enrichment score and FDR, false discovery rate. (C) Top 15 enriched Gene ontology (GO) biological processes (BP) by GSEA in gRNA mix (denoted in red) and no gRNA (denoted in green). Vertical axis represents GOBP terms, and horizontal axis represents normalized enrichment score (NES). (D) GSEA plot of genes involved in immune system development, negative regulation of cell cycle and DNA damage in dCas9-KRAB with gRNA mix compared to no gRNA cells.

The silencing of *EWSR1-FLI1* in A673 cells transduced with dCas9-KRAB with gRNA mix differentially regulated a large set of 4557 (1863 downregulated and 2694 upregulated genes) and 6408 annotated genes (3070 downregulated and 3338 upregulated genes) relative to dCas9-KRAB with no gRNA and non-transduced A673, respectively ((|log_2_(FC)| > 1, *p*-adj < 0.05). Despite slight non-specific differences in gene expression, transcriptome profiling revealed high correlations between dCas9-KRAB with no gRNA and non-transduced cells (Figure 3A), consistent with Thakore *et al*.,^24^ which reported slight changes in gene expression between non-transduced cells and cells transduced with dCas9-KRAB with gRNA or no gRNA that could be attributed to the viral transduction process.

Subsequently, RNA-seq of *EWSR1-FLI1* silenced A673 cells (dCas9-KRAB gRNA mix) and the control (no gRNA) cells were compared against published EWS-FLI1 target gene signatures.^67,68^ Gene set enrichment analysis (GSEA) revealed that the reported EWSR1-FLI1 down-regulated genes^68^ were derepressed by the epigenetic silencing of *EWSR1*-*FLI1* (gRNA mix) in A673 (Figure 3B, top; and Supplemental Table 1). Likewise, the reported EWSR1-FLI1 up-regulated genes were depleted in the *EWSR1*-*FLI1* silenced cells (gRNA mix) compared to the no gRNA control (Figure 3B, top). GSEA of another EWS-FLI1 gene signature obtained from ectopically expressed EWS-FLI1 in human MSCs^67^ also showed similar enrichment patterns (Figure 3B, bottom), confirming that dCas9-KRAB gRNA mix can reverse the gene expression profiles induced by *EWSR1-FLI1* in A673 cells.

As expected from previous work, our Gene Ontology (GO) analysis revealed that the derepressed genes in the gRNA mix were mainly involved in immune function^69^ (such as immunoglobulin production, immune system development, and B cell activation) and other processes, such as cellular response to UV, response to DNA damage signaling,^70^ negative regulation of the cell cycle^71^ and protein localization to the nucleus (Figure 3C, top; and Supplemental Table 2). Specifically, we observed upregulation of genes encoding various cytokines and cytokine receptors (e.g., C–X–C motif chemokine 12 (*CXCL12*), Interleukins and their receptors (*IL-15*, *IL7R*, *IL13R2*), and the tumor necrosis factor superfamily member 4 (TNFSF4)), immune checkpoint blockade (*CD274*/*PD-L1*), antigen processing and presentation targets (e.g., major histocompatibility complex gene (*HLA-B*), transporter associated with antigen processing (*TAP1*, *TAP2*), and endoplasmic reticulum aminopeptidase 2 (*ERAP2*)), DNA repair gene (poly(ADP-ribose) polymerase 9 (*PARP9*)) and key regulator of the cell cycle progression (cyclin-dependent kinase (*CDK6*)) associated with downregulation of Cyclin D1 (*CCND1*)).^71–74^ The enriched genes in the no gRNA control (i.e., upregulated in the presence of *EWSR1-FLI1*) cells revealed significant involvement in nervous system development, including neurotransmitter transporter and secretion, neuromuscular process, regulation of dendrite development, and regulation of vesicular neurotransmitter transporter (Figure 3C, bottom; and Supplemental Table 2 and 3).

In summary, these transcriptional analyses provided support that our epigenetic editing approach was mediated by *EWSR1*-*FLI1* gene repression and it was consistent with the *in vitro* phenotypes associated with changes in proliferation and migration produced by dCas9-KRAB.

### *WSR1-FLI1* silencing by dCas9-KRAB decreased tumor growth in EWS xenograft mouse models

We next assessed whether the dCas9-KRAB gRNA mix targeting *EWSR1-FLI1* could inhibit the growth of A673 cells *in vivo.* To this aim, 4-week-old female NOD-SCID-IL2 receptor gamma null (NSG) immunocompromised mice were randomly assigned into three groups (*n = 36*, 12 mice per group). Mice were injected subcutaneously into the right flank of NSG mice with either 1) A673 cells lentivirally transduced with dCas9-KRAB and gRNA mix, 2) A673 cells transduced with dCas9-KRAB with no gRNA, or 3) A673 non-transduced cells. After implantation, tumor size was monitored continuously during the experiment (Figure 4A).

**Figure 4.**
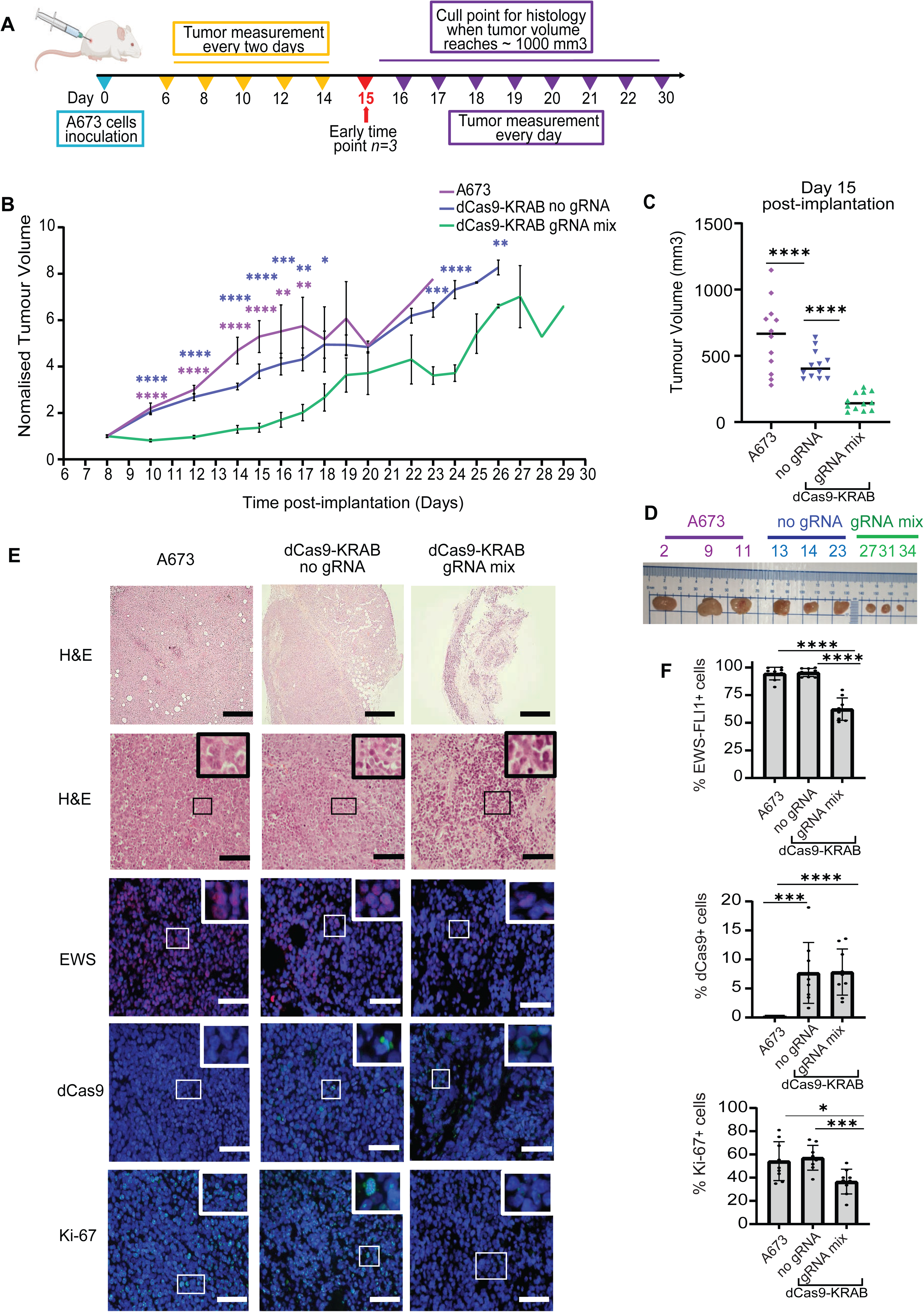
Suppression of *EWSR1-FLI1* in EWS A673 transduced xenograft models by dCas9-KRAB. (A) Schematic representation of experimental timeline. A673 cells transduced with either dCas9-KRAB gRNA mix, dCas9-KRAB no gRNA, or A673 non-transduced cells were resuspended in a 1:3 mixture of Matrigel and ice-cold serum-free DMEM media and inoculated subcutaneously into NSG mice at day 0 (*n = 12* per group). Tumor volumes were monitored every two days using a digital Vernier. Animals were sacrificed at day 15 for histological examination (n = 3 per group). Mice were euthanized at the experimental endpoint for histological examination (when tumor volumes < 1000 mm^3^). Data are plotted as mean values ± SEM and statistical significance was determined using multiple unpaired *t*-test (\**p* < 0.05, \*\**p* < 0.01, \*\*\**p* < 0.001, \*\*\*\**p* < 0.0001). (B) Average tumor volume measurements throughout the experiment normalized to day 8. (C) Scatter plot indicating the tumor volume (mm^3^) of each mouse at day 15 post-implantation. Data are plotted as mean values ± SEM and statistical significance was determined using multiple unpaired *t*-test (\*\*\*\**p* < 0.0001). (D) Representative images of tumors extracted at day 15 post-inoculation of the tumor cells. (E) Representative sections of tumors collected at day 15 from (n= 3 mice per group) stained with hematoxylin and eosin (H&E), and for the detection of the proliferation marker (Ki-67) (green), EWS-FLI1 (red), and dCas9 (green) antibodies. (F) Quantification of these markers is shown on the right panels. Data are plotted as mean values ± SD and statistical analysis was carried out using multiple unpaired *t*-test (\*\**p* < 0.01, \*\*\**p* < 0.001, \*\*\*\**p* < 0.0001). Scale bars represent 50 µm.

Notably, from day 1 to 15 post-implantation of the cells, we detected a significant reduction in tumor growth in A673 xenografts expressing dCas9-KRAB-gRNA mix by more than 50%, (tumor size average = 155 mm^3^ at day 15) as compared to the dCas9-KRAB no gRNA group (tumor size average = 432 mm^3^, *p* < 0.0001) or A673 non-transduced control (tumor size average = 601 mm^3^, *p* < 0.0001) (Figure 4B). From day 16 to day 29 post-implantation, an increase in tumor growth was observed for the gRNA mix group; however, sizes were still significantly smaller than the controls, even at experimental end point. Additionally, no significant changes were observed in the tumor volume between A673 expressing dCas9-KRAB no gRNA group and the non-transduced control group during the experiment, except for day 14, where the average tumor volume was 357 mm^3^ and 532 mm^3^ for no gRNA and A673 groups, respectively, *p* = 0.017. (Figure 4B). Once the A673 control tumors reached the experimental endpoint of approximately 1000 mm^3^, three randomly selected mice per group were humanely euthanized, and resected tumor tissues were processed for histopathological evaluation (Figure 4C,D).

Consistent with decreased tumor growth, the histological assessment by IF (Figure 4E,F) at day 15 post-implantation of the cells showed significant EWS-FLI1 protein downregulation in A673 transduced with dCas9-KRAB-gRNA mix (frequency of EWS-FLI1^+^ cells = 62%, *p* < 0.0001) compared to the control groups (non-transduced A673 and no gRNA groups, 95% and 94%, respectively). Remarkably, the expression of dCas9-KRAB protein was detected in 8% of cells in both gRNA mix and no gRNA groups, but as expected, it was undetected in non-transduced A673 cells. The relatively small percentage of dCas9-KRAB positive cells in the resected tumors suggests a possible negative clonal selection of these transduced cells during tumor progression in the dCas9-KRAB gRNA mix. This is in accordance with the relative recurrence of the tumors after day 15, and the loss of repressive capacity *in vivo* as compared to that of the cell line prior to implantation (Figure 1G). Consistent with this notion, IF assessment of tissue sections obtained from mice at late time point demonstrated a loss of dCas9-KRAB expression relative to early time points. Late time point tumors regained the expression of EWS-FLI1, which was similar to that of non-transduced control groups (Figure S7, Supporting Information 1).

As expected from the tumor growth curves above, a significant reduction of tumor cell proliferation index (Ki-67) was observed in A673 resected tumors transduced with dCas9-KRAB with gRNA mix (36%, *p* < 0.001) compared to no gRNA (56%) and non-transduced control (54%) groups (Figure 4E,F). However, no significant changes between the examined groups were noticed in expressing the apoptotic marker (cleaved caspase-3), indicating that the decreased growth in dCas9-KRAB with gRNA mix tumors was most likely attributed to reduced tumor cell proliferation rather than an increase in apoptosis (Figure S8, Supporting Information 1).

### Repression of *EWSR1-FLI1* using CRISPR/RNPs delivered with engineered polymeric nanoparticles

While gRNA combinations co-delivered with dCas9-KRAB protein showed strong silencing of *EWSR1-FLI1*, these proof-of-concept studies were performed with lentiviral vectors which are not suitable for clinical applications. Therefore, we next investigated non-viral delivery systems to transduce purified dCas9-KRAB protein in combination with synthetic gRNAs (referred to as ribonucleoprotein complexes, RNPs) to induce specific *EWSR1-FLI1* oncogenic silencing. For non-viral delivery, we utilized a polymeric dendritic formulation, a generation 5 (G5) dendronized polymer partially functionalized with *p*-guanidinobenzoic acid (a GBA-polymer), previously shown by our laboratories to effectively deliver protein cargoes into cancer cells (Figure 5A).^75^

**Figure 5.**
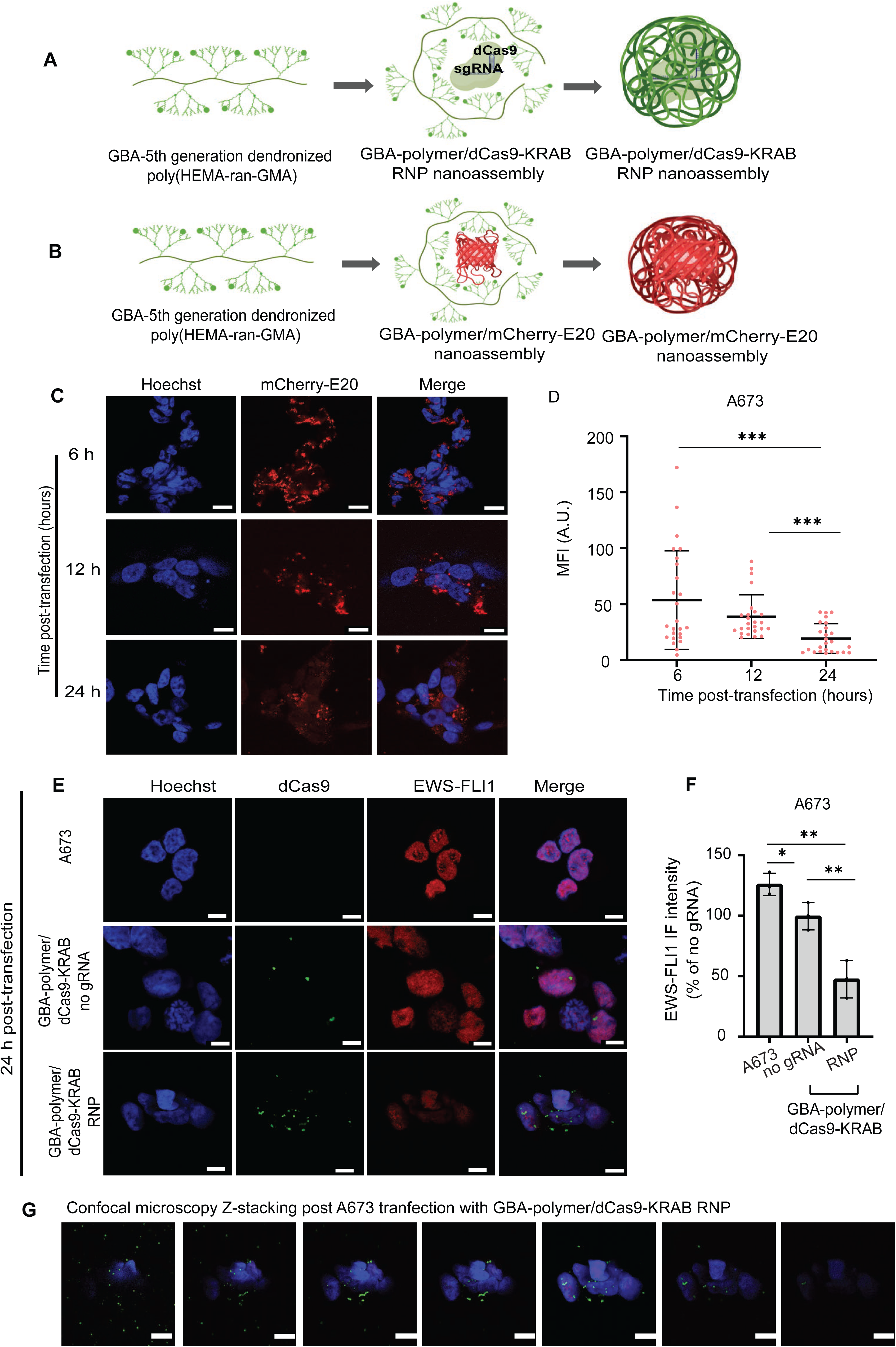
Generation of a GBA-polymer nanocarrier to repress *EWSR1-FL1* in A673 cells. (A,B) Schematic representation of hierarchical nanoassemblies between the reporter mCherry-E_20_ protein (A) and dCas9-KRAB (B) with GBA-polymers associated via electrostatic interactions. C) Representative images for intracellular delivery of mCherry-E_20_ protein by PAMAM-polymeric nanoparticles in A673 cells at mass ratio 1.5:1 (GBA-polymer: mCherry-E_20_) after 6, 12, and 24 h of transfection; mCherry-E_20_ (red) and nuclei (Hoechst, blue). Quantification of mCherry-E_20_ protein is shown on the right panel (D). Data are presented as mean values ± SD. *p*-values were determined by Brown-Forsythe and Welch ANOVA tests with Dunnett’s T3 multiple comparison test (\*\**p* < 0.01, \*\*\**p* < 0.001). E) Representative images for dCas9-KRAB protein delivery into A673 cells using the PAMAM- polymeric nanocarrier at mass ratio 1.5:1 (polymer: protein) after 24 h post-transfection. EWS-FLI immunostaining (red), dCas9 immunostaining (green) and nuclear Hoechst staining (blue). (F) Mean fluorescent intensity (MFI) of EWS-FLI1 protein after 24 h of transfection by polymer. Data are normalized to the dCas9-KRAB no gRNA and presented as mean values ± SD. *p*-values are determined by multiple unpaired *t*-test (\**p* < 0.05, \*\**p* < 0.01). *n = 3* biologically independent experiments. Scale bar represents 10 μm. G) Representative images of confocal microscopy Z-stacking for the cytoplasmic and nuclear delivery of dCas9 protein using polymeric system, 24 h post-transduction of the cells. Z-stacking was carried at 100 nm intervals.

We first produced *Sp*dCas9-KRAB protein from *E. coli* using a Maltose Binding Protein (MBP) recombinant vector system, in which dCas9-KRAB was C-terminally fused to the MBP carrier. The construct was engineered with three Nuclear Localization Sequences (NLS), three FLAG tags to enable the detection of the protein in mammalian cells, and with poly-glutamic sequence comprising 20 residues to provide a negatively charged domain to the protein to facilitate binding to our positively charged polymer (Figure S9A, Supporting Information 1).^76^ After enzymatic cleavage of the MBP, we purified NLS-FLAG-Etag-dCas9-KRAB-NLS protein (abbreviated as dCas9-KRAB) nearly at 75% purity (Figures S9B,C, Supporting Information 1).

For intracellular delivery experiments facilitated by the GBA-polymer, we first tested the delivery of reporter protein mCherry linked to an array of 20 glutamic acids (mCherry-E_20_). mCherry-E_20_ was harnessed as a positive control to monitor cellular uptake and protein cytosolic delivery into the cells by real-time immunofluorescence in A673 cells (Figure 5B). Efficient delivery of the reporter protein in this system was validated from 6 h to 24 h post-transfection, where the maximum intensity of mCherry-E_20_ was observed at 6 h in accordance with previous work in multiple other cell types (Figure 5C,D).^37^

We next examined the efficacy of the polymeric platform in delivering dCas9-KRAB in complex with the three synthetic gRNAs into the same cells. Since it has been shown that Cas9/RNPs achieve maximum genome editing efficacy at 24 h-post transfection,^35,77^ the RNP/GBA-polymer nanoassemblies were similarly incubated with A673 cells for 24 h to enable the assessment of intracellular delivery and gene regulation (Figure 5E). IF analysis revealed significant repression of EWS-FLI1 protein by GBA-polymer/dCas9-KRAB in presence of gRNAs (hereafter referred to as dCas9-KRAB RNP) (48%, *p* < 0.01) relative to GBA-polymer/dCas9 with no gRNA (hereafter referred to as dCas9-KRAB no gRNA) and A673 control (Figure 5F). The intracellular localization of the dCas9-KRAB in the absence or presence of gRNAs was confirmed by confocal microscopy using an anti-dCas9 antibody (Figure 5G). Lastly, the effects of dCas9-KRAB RNP in repressing EWS-FLI1 were attenuated beyond 24 h post-transfection to (78%, *p* = 0.03) relative to dCas9-KRAB no gRNA at 72 h post-transfection (Figure S10, Supporting Information 1), consistent with the decay of the expression of dCas9-KRAB (Figure 5A).

### CRISPR/RNPs assembled with engineered polymeric nanoparticles silence *EWSR1-FLI1 in vivo* and inhibit tumor growth in mouse models of EWS

We subsequently evaluated the capacity of the GBA-polymeric delivery system for dCas9-KRAB/RNP targeting *EWSR1-FLI1* to inhibit tumor growth in mouse models of EWS. First, we generated subcutaneous xenografts by injecting 1.5 × 10^6^ A673 cells into the right flank of 4-week-old NSG mice. To assess the expression and stability of RNPs i*n vivo*, when tumors reached 150-200 mm^3^, mice were either injected orthotopically with a single dose of active dCas9-KRAB RNPs (with the active mix of three gRNAs targeting *EWSR1-FLI1*), and dCas9-no gRNA, or PBS (vehicle control), with *n=12* mice/group. Animals were sacrificed (*n=3* mice/group) at 4, 8, 12, and 24 h post-intratumoral injection of the formulations for IF assessment of dCas9-KRAB and EWS-FLI1 expression. Tumor biopsies collected at 4 and 8 h from mice treated with dCas9-KRAB no gRNA and dCas9-KRAB RNP showed higher expression of dCas9-KRAB protein compared to 12 and 24 h post-injection (Figure 6A). Importantly, dCas9-KRAB RNP decreased the expression levels of EWS-FLI1 at 4 h post-injection relative to dCas9-KRAB no gRNA (Figure 6B).

**Figure 6.**
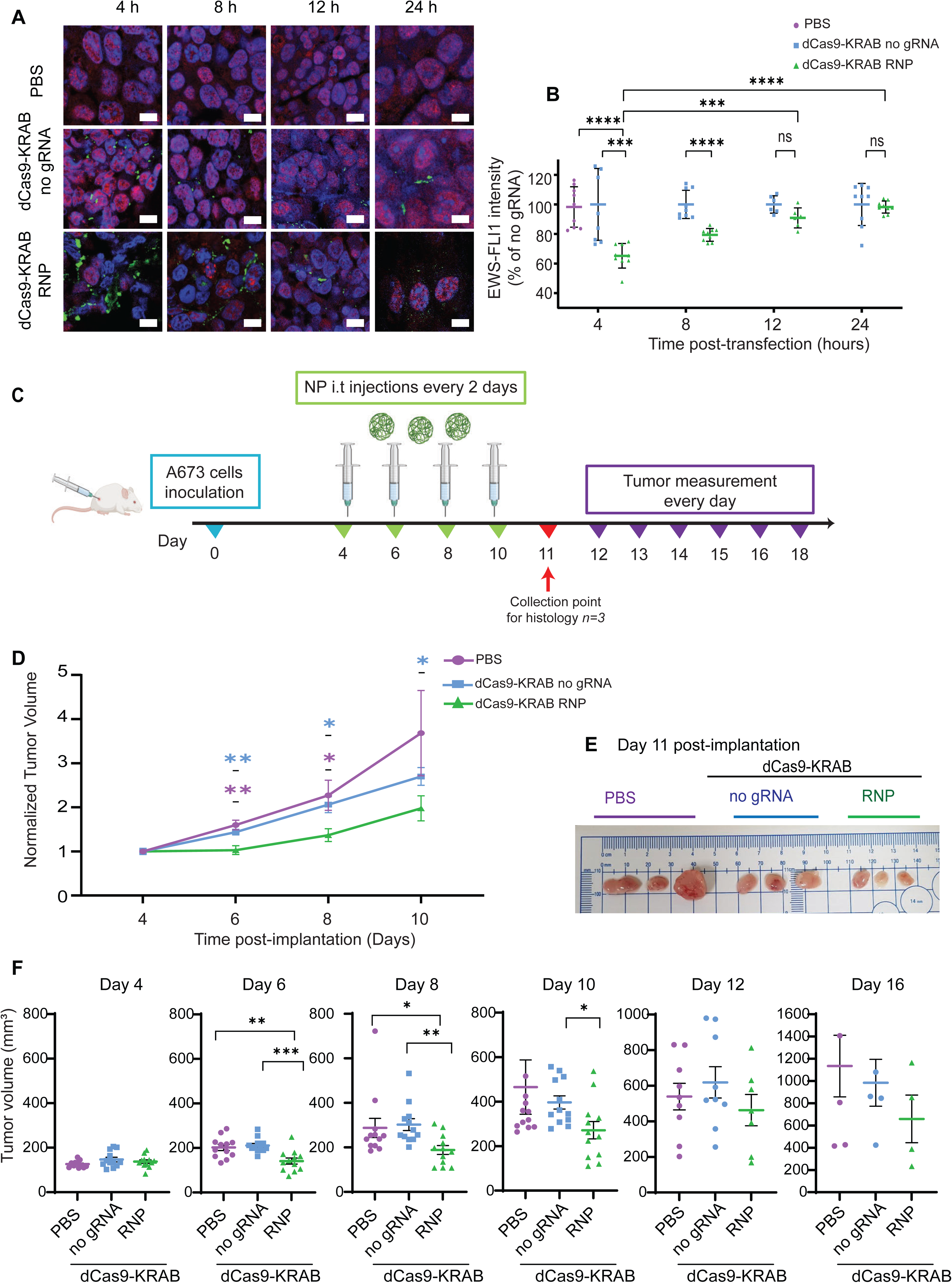
Repression of the oncogenic fusion *EWSR1-FLI1 in vivo* by dCas9-KRAB/RNP delivered by the GBA-polymeric system. (A) Representative sections of tumors collected at 4, 8, 12, 24 h post-last intratumoral injection (3 mice per group) and stained with anti-CRISPR/Cas9 (green) antibodies and anti-EWS-FLI1 (red). Scale bars represent 20 µm. (B) Quantification of the two markers is shown on the right panels. Data are plotted as mean values ± SD and statistical analysis was carried out using two-way ANOVA with Tukey’s multiple comparison tests (\*\*\**p* < 0.001, \*\*\*\**p* < 0.0001). Scale bars represent 20 µm. (C) Schematic timeline of the experiment. 1.5 × 10^6^ cells/100 µL A673 tumor cells were inoculated subcutaneously (day 0) into NSG mice (N = 36 mice). Mice received a total of four intratumoral injections between day 4 and day 10, one injection every 48 h (*n = 12* mice per group). Mice were humanly sacrificed 24 h-post last injection for histological analysis (*n = 3* per group). Control animals were culled after day 12, when reached 1000 mm^3^. (D) Relative tumor volume in A673 xenografts treated intratumorally with dCas9-KRAB RNP, dCas9-no gRNA, or PBS. Tumor volumes were normalized to day 4 post-implantation of the cells. Representative images of A673 resected tumors harvested at day 11 are indicated in (E). Data are presented as mean values ± SEM (*n = 12* per group) and statistical analysis was undertaken using multiple unpaired *t*-test (\**p* < 0.05, \*\**p* < 0.01). (F) Relative changes in tumor volumes of individual mice at days 4, 6, 8, 10, 12 and 16 post-implantation. Data are plotted as mean values ± SEM (*n = 12* per group) and statistical analysis was undertaken using multiple unpaired *t-*test (\**p* < 0.05, \*\**p* < 0.01, \*\*\**p* < 0.001).

To assess therapeutic efficacy, the same formulations as above were injected subcutaneously into A673 xenografts (*n = 36* NSG mice; Figure 6C). When tumor volumes reached an average size of ∼150 mm^3^, mice were randomized into three groups (*n = 12* mice/group) and injected intratumorally with either dCas9-KRAB RNP, dCas9-KRAB no gRNA or with vehicle control (PBS). Animals were administered a total of four injections (one injection every 2 days), starting at day 4 post-implantation of the cells. Tumor growth was monitored every two days and normalized to that of day 4 prior to treatment. At day 4, no significant changes were observed in tumor volumes between the groups. In contrast, at day 6, there was a significant 33% and 30% reduction in tumor volumes for the dCas9-KRAB RNP group compared to dCas9-KRAB no gRNA and PBS groups, respectively (vehicle: 202 mm^3^, *p* = 0.0045; no gRNA group: 211 mm^3^, *p* = 0.0003, dCas9-KRAB RNP: 141 mm^3^). By day 8, the average tumor volumes were significantly 1.6-fold lower for the dCas9-KRAB RNP group compared to dCas9-KRAB no gRNA treatment group (188 mm^3^ and 302 mm^3^ respectively, *p* = 0.0023) and 1.5-fold lower for the mice receiving dCas9-KRAB RNP relative to PBS group (287 mm^3^, *p* = 0.0483). At day 10, dCas9-KRAB RNP sustained a significant antitumoral effect, with an average tumor size of 271 mm^3^ relative to dCas9-KRAB no gRNA treatment group (396 mm^3^, *p* = 0.018) (Figure 6D,F). However, no significant changes were observed in tumor volumes between groups at day 12 and day 16 (Figure 6F).

Importantly, no significant changes in tumor volumes were detected in a similar experiment in which A673 mouse xenografts were injected with dCas9-KRAB RNP in the presence of heterologous/nonspecific gRNAs designed against a control gene that is not expressed in mammalian cells (the *GFP* gene). This experiment demonstrates that the reduction in tumor growth was dependent on the *EWSR1-FLI1* specific gRNAs (Figure S11, Supporting Information 1).

### CRISPR/RNPs assembled with engineered polymeric nanoparticles silence *EWSR1-FLI1* in PDX mouse models of EWS

To validate our formulation into a more translational setting, we interrogated tumor growth inhibition in a patient-derived xenograft of Ewing Sarcoma (ES-4).^78–80^ Confirmatory diagnosis of EWS for ES-4 tumor was performed by histopathology, immunohistochemistry, and molecular cytogenetics (fluorescence *in situ* hybridization (FISH) analysis). Hematoxylin and eosin (H&E) staining showed typical histomorphology defined by the presence of monomorphic small round blue cells with abundant clear cytoplasm, and fine, stippled chromatin patterns. Immunohistochemical examination revealed cells showing strong and diffuse positive membranous protein CD99, and sections were also positive for the EWS marker. Finally, FISH assays confirmed the predicted rearrangement of the *EWSR1* gene at chromosome 22q12 and the *EWSR1-FLI1* rearrangement at the abnormal chromosome 22 (Figure S12, Supporting Information 1).

The ES-4 PDX tumor tissues were surgically implanted subcutaneously and successfully grown in a total of 20 mice for intratumoral injection of polymeric formulation regimes. When tumors reached approximately 100 mm^3^, mice were randomized into two groups (n = 10 mice/group) and received intratumoral injections of either dCas9-KRAB RNP or dCas9-KRAB no gRNA. Animals were given a total of four injections (one injection every 2 days, Figure 7A). Tumor growth was measured every 2 days with volumes normalized immediately prior to the first treatment.

**Figure 7.**
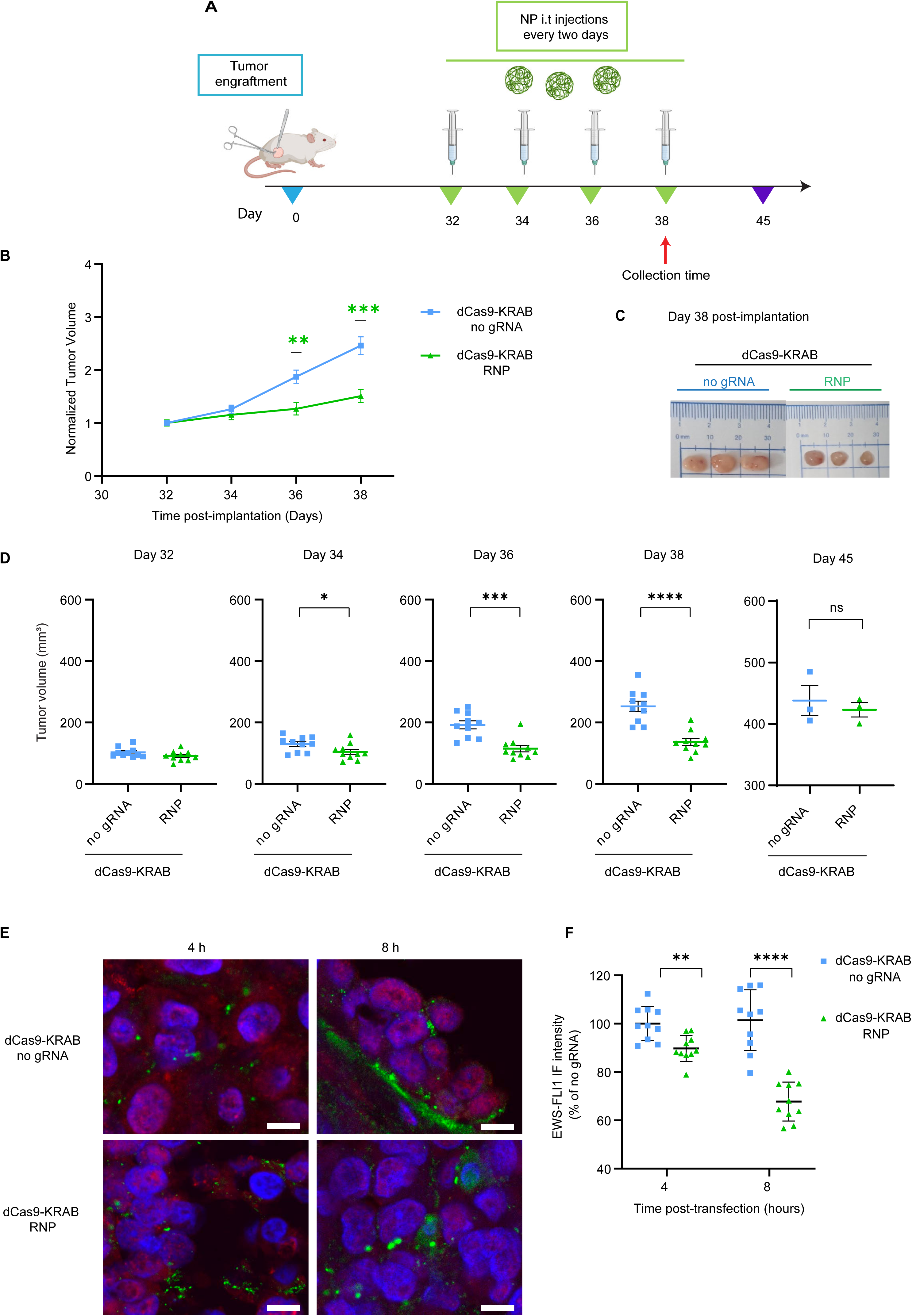
Repression of the oncogenic fusion *EWSR1-FLI1* in PDX mouse models of EWS by dCas9-KRAB/RNP delivered by the GBA-polymeric system. (A) Experimental timeline outlining treatment regimes in ES-4 PDXs: mice were treated every 48 h, receiving a total of 4 intratumoral injections. (B) Mean tumor volume measurements in EWS PDX mice injected intratumorally with dCas9-KRAB RNP or dCas9-KRAB no gRNA control. Tumor volumes are normalized to their sizes prior to the first treatment. Data are plotted as mean values ± SEM (n = 10 per group). Statistical significance was performed using multiple unpaired *t*-test (\*\**p* < 0.01 and \*\*\**p* < 0.001). (C) Representative images of the resected tumors at 4 h post-last injection (n = 3 per group). (D) Volumetric scatter plots of individual tumors at days 32, 34, 36, 38 and 45 post-implantation. Statistical significance was performed using multiple unpaired *t*-test (\**p* < 0.05, \*\*\**p* < 0.001 and \*\*\*\**p* < 0.0001). (E) Representative sections of tumors collected at 4 and 8 h post-last intratumoral injection (3 mice per group) and stained with anti-CRISPR/Cas9 (green) antibodies and anti-EWS-FLI1 (red). (F) Quantification of these markers is shown on the right panels. Data are plotted as mean values ± SEM and statistical analysis was carried out using two-way ANOVA with Tukey’s multiple comparison tests (\**p* < 0.05, \*\*\*\**p* < 0.0001). Scale bars represent 20 µm.

Prior to the second dose, we observed a rapid and significant reduction of tumor growth (*p* = 0.041) (day 34 post-implantation) between dCas9-KRAB RNP-treated mice (with EWS gRNA mix) and the dCas9-KRAB no gRNA group (average tumor growth of dCas9-KRAB RNP: 104 mm^3;^ dCas9-KRAB no gRNA group: 130 mm^3^). Prior to the third dose (day 36 post-implantation) it was a 40% reduction in tumor growth promoted by dCas9-KRAB RNP (dCas9-KRAB RNP: 115 mm^3^; no gRNA control: 192 mm^3^, *p* = 0.0002). Lastly, prior to the fourth dose (day 38 post-implantation), the dCas9-KRAB RNP (with EWS gRNA mix) treatment group still exhibited a significant therapeutic effect relative to no gRNA with an average tumor size of 136 mm^3^ and 252 mm^3^ (*p* < 0.0001), respectively (Figure 7B,D). Tumors were collected at 4 h and 8 h post-last injection and tumor biopsies were examined histologically by IF. We observed high expression of the dCas9-KRAB protein in the resected tumors, which was associated with a significant reduction in the expression of EWS-FLI1 by 10% and 33% at 4 h and 8 h post-injection, respectively, relative to no gRNA (Figure 7E,F). However, twelve days after the last dose (day 45), at day 45, no significant differences in tumor volumes between the groups were observed (Figure 7D).

In conclusion, our results show that dCas9-KRAB/gRNAs could be successfully delivered using dendritic polymers. Despite their transient “hit-and-run” and transient expression in the tumor cells, we show that these formulations were able to inhibit *EWSR1-FLI1* oncogenic expression *in vivo* and quickly decrease/debulk tumor growth post-injection, with an effect lasting at least 12 days post-orthotopic treatment of the tumors.

## Discussion

This manuscript outlines for the first time a polymeric formulation that promotes efficient delivery of dCas9-KRAB RNP complexes when injected in EWS tumors, causing a fast and significant tumor inhibition in highly aggressive models of EWS disease. EWS remains a highly aggressive pediatric malignancy. The quality of life and outcomes for patients with metastatic or relapsed tumors remains extremely poor, with a 5-year survival rate of <30%.^47,81^. The EWS-FLI1 is the main driver oncogene in EWS and its continuous expression is essential for tumorigenesis; we therefore reasoned that epigenetic repression of such cancer specific EWS-FLI1 fusion represents an excellent model system to assess the efficacity of dCas9-KRAB RNP delivery in impacting tumorigenesis *in vivo*.

We identified a combination of three gRNAs that significantly silenced the expression of *EWSR1-FLI1* at the mRNA and protein levels in CRISPR-treated EWS cells and to a greater extent than those of each individual gRNAs. Of note, previous studies demonstrated a degree of efficacy of small interfering RNAs (siRNAs) and shRNA systems in repressing *EWSR-FLI1*; however, the significant off-target activities of these technologies represent a serious concern.^68,82–89^ On the other hand, while highly selective CRISPR/Cas9 systems cause irreversible gene disruptions, the dCas9-KRAB system does not alter the underlying DNA sequence but silences endogenous genes by promoting chromatin compaction, minimizing the potential of genomic rearrangements and off-target activity post-treatment.^89–91^ Interestingly, suppressing oncogenic fusions by the CRISPR/dCas9 system requires targeting the regulatory regions of the first fused genes without designing new gRNAs, siRNA or shRNA whenever the second fused gene or isoform type is different.^90,92–96^ In contrast with more complex arrays of effectors, we found that the RNA-guided dCas9-KRAB displayed negligible off-target gene expression, consistent with previous studies.^13,24,57,97–102^ In addition, KRAB, as a single potent repressive effector domain, has a small and negatively charged structure, which is also expected to facilitate encapsulation in positively charged vehicles.^103–109^ In contrast with short-lived RNAi approaches, our laboratory and others have shown long-lasting therapeutic effects with dCas9 fused with epigenetic silencing domains, particularly with KRAB promoting strong chromatin condensation and gene silencing *in vivo*.^61,110^ In fact, in a recent study, we show that dCas9-KRAB has the potential to induce long-lasting genome-wide epigenetic changes in triple-negative breast cancer models.^61^ Therefore, while herein we utilized EWS tumor models, the tools described in this manuscript have a broad range of applications to silence multiple other oncogenic drivers in other solid tumors.

As expected from the *EWSR1-FLI1* silencing promoted by dCas9-KRAB, we observed strong inhibition of the tumorigenic phenotype (with virtually no colonies formed in semisolid media), inducing significant antiproliferative effects and strong inhibition of EWS cancer cell migration. Similar to previous studies, our findings add supporting evidence to the notion that Ewing sarcoma cells are *EWSR1-FLI1*-dependent.^65,86,91,111^ In our hands, epigenetic editing of the wild type EWSR1 by dCas9-KRAB resulted in transcriptional downregulation, although the endogenous expression of the EWSR1 promoter in normal cells (e.g. in primary HDFa) is significantly lower than that in EWS cells. We did not observe significant changes in cell proliferation induced by dCas9-KRAB in normal HDFa cells, suggesting that normal cells are less dependent or addicted to EWSR1 promoter activity than EWS cells.

Our Analyses of RNA sequencing data identified over 4000 genes that were differentially regulated in the A673 cells treated with dCas9-KRAB with gRNA mix, reinforcing the notion that EWS-FLI1 TF reprograms the transcriptional landscape of EWS cells.^112^ Interestingly, the number of upregulated genes (2694) was higher than downregulated ones (1863) post-*EWSR1-FLI1* suppression, indicating that gene repression promoted by EWS-FLI1 may be more prevalent than transcriptional activation in A673 cells, consistent with previous works.^85,113–115^ Additionally, our DEG set induced by dCas9-KRAB was positively correlated with DEGs modulating *EWSR1-FLI1* with other methods (shRNA knockdown and ectopic cDNA delivery of the oncogenic fusion)^67,68^ demonstrating that our dCas9-KRAB regulated known *EWSR1-FLI1-*dependent target genes.

Examination of the transcriptome of the dCas9-KRAB edited cells offers an opportunity to identify therapeutic weaknesses that could be exploited in conjunction with our RNP approaches to improve phenotypic outcomes. Interestingly, our GO analysis revealed that dCas9-KRAB transduced with gRNA mix in A673 cells upregulated genes involved in immune-related processes. These genes include multiple cytokines and cytokine receptors, such as *CXCL12*, *IL-15*, *IL7R*, *IL13R2*, and *TNFSF4*. Importantly, we also observed upregulation of the immune checkpoint blockade *CD274/PD-L1*, together with significant upregulation of some antigen processing and presentation targets, such as *HLA-B*, *TAP1*, *TAP2*, and *ERAP2*. This is consistent with previous work in A673 cells, showing that EWS-FLI1 suppression by shRNA similarly induced upregulation of IL-6 and other cytokines such as *CXCL1* and *GM-CSF* ^69^. Therefore, it is plausible that silencing this fusion could reprogram the immune response and that our RNP approach could be used in the future in conjunction with immune-based therapies. Our GO analysis also revealed enrichment of upregulated genes in cellular response to UV and DNA damage, such as *PARP9*, *TREX1*, and *ERCC6*. Consistently, recent reports also reported that EWS depletion results in alternative splicing changes in genes involved in DNA repair and genotoxic stress signaling,^70^ and that EWS cells display alterations in regulation of damage-induced transcription.^116^ These results suggest that combination of our RNPs with drugs targeting the DNA repair pathways (such as PARP inhibitors) or cell cycle progression (such as CDK6 inhibitors) could increase the efficacy of dCas9-KRAB with gRNA mix system.

In terms of the longevity of the anti-tumor phenotype, we reported a potent antitumor response with dCas9-KRAB with gRNA mix in EWS cell-derived xenografts mouse models for a period up to approximately 15 days post-implantation of the cells when using lentiviral vehicles and stably transduced cell lines. However, we reported a progressive loss of the expression of dCas9-KRAB after day 15, which was concurrent with the re-expression of *EWSR1-FLI1* in the tumor cells over time, both observations consistent with tumor recurrence. Similar to Takigami *et al*., the transient reduction in tumor growth could be attributed to significant repression of the proliferative activity of the treated EWS cells rather than induced apoptosis.^117^ Remarkably, previous work that utilized RNA interference (RNAi) or CRISPR (nuclease competent) methods also revealed significant tumor growth reduction, although with partial loss of RNAi through the course of the study.^65,86,91^ To assess this limitation, multiple cycles of treatments involving CRISPR/dCas9 deliver potentially in combination with chemotherapies, therapies targeting DNA damage, and immune-base therapies in immune-competent animals should be envisaged in the near future.

Our functional proof-of-concept studies to deliver dCas9-KRAB and gRNAs were first performed using lentiviral vectors. However, lentiviruses or even adenoviral vehicles to deliver the CRISPR biomolecular components as episomal plasmid payload/s suffer from several drawbacks, such as the possibility of insertional mutagenesis and immunogenicity.^93^ Seminal studies applied CRISPR/Cas9-based strategy delivered by lentiviral or adenoviral systems to successfully target oncogenic fusions, including *BCR-ABL*,^91,92^ *PAX3-FOXO1*,^94^ *TMEM135-CCDC67* and *MAN2A1-FER*.^95^ Cervera *et al*. utilized lentiviral vectors to deliver CRISPR components, in the form of pDNA, to inactivate *EWSR1-FLI1* by targeting exon 9 of the *FL1* gene in A673 cells, resulting in blockage of cell proliferation and induction of cellular senescence.^90^ Further, Martinez-Lage *et al*. used lentiviral and adenoviral vectors to deliver CRISPR/pDNA elements *in vitro* and *in vivo*, respectively, to induce permanent inactivation of *EWSR1-FLI1* by targeting the intronic regions of *EWSR1* and *FLI1* genes, producing significant reduction of cellular proliferation and clonogenicity, and tumor growth inhibition.^91^ Nevertheless, despite the therapeutic potential of the CRISPR/Cas9 system for fusion-driven cancers described in these studies, there were shortcomings associated with viral delivery systems that restricted their clinical application.

The delivery of CRISPR as RNPs offers an alternative approach over plasmid vectors, particularly for epigenetic editing approaches, which can benefit remarkably from transient “hit and run” delivery methods yet elicit potentially durable responses.^118,119^ Furthermore, there is evidence that RNP delivery, while transient, results in high on-target specificity, lower off-target effects, and lower immunogenicity than plasmid-encoded transduction systems.^118,119^ However, the large size of dCas9 protein (∼160 kDa) and poor endosomal escape are the main obstacles that limit the application of the RNP approach.^120,121^ While several nanocarriers have been described for the delivery of Cas9/RNPs,^41–43,118,120,122,123^ only a limited number of nanoparticles have been applied for the encapsulation of dCas9-effector domain/RNPs including lipid nanoparticles (LNPs),^124^ polymer-based nanoparticles,^125–127^ and extracellular vesicles.^128^ To the best of our knowledge, we provide the first report showing *in vivo* delivery of dCas9-KRAB RNPs in xenografts and PDXs.

Compared with lipid vehicles, cationic polymers were found to be one of the most promising alternatives for CRISPR/RNP *in vivo* delivery, demonstrating high stability and biocompatibility, lower immunogenicity, less restriction in payload size, simplicity of synthesis and functionalization, greater endosomal escape, and controlled-release chemical structure.^129–134^ Lee *et al.* used magnetic peptide-imprinted polymers to effectively deliver the targeted CRISPR/dCas9-activation system in the form of RNP into HEK293T cells, which achieved highly efficient activation of OSKM,^125^ INS,^126^ and SNCA^127^ genes for cellular reprogramming,^125^ upregulation of insulin expression^126^ or for generating an *in vitro* model for the aggregation of the SNCA protein in neurodegenerative diseases.^127^ Recently, RNPs encapsulating dCas9-VPR activators have been delivered *in vitro* for the treatment of glioblastoma multiforme^135^. However, further validation of these strategies for animal models is yet to be undertaken.

Here, we harnessed polymeric formulations to deliver RNP payloads where dCas9 has been modified with a negatively charged poly(glutamic acid) (E_20_) tag to enhance the encapsulation of RNPs.^136^ The polymer utilized in this study has previously been shown to deliver the reporter control protein mCherry in different cell lines with high efficiency and low toxicity while minimizing endosomal degradation, one of the main obstacles to intracellular delivery of cargos.^37,44,75^ Further, our findings demonstrated a very significant reduction in *EWSR1-FLI1* expression *in vitro* mediated by the polymer in A673. In *in vivo* experiments, we found that dCas9-KRAB had the highest expression in the injected tumors at 4 h and 8 h post-injection, with the expression persisting even at 24 h post-injection, when we observed a strong reduction of *EWSR1-FLI1* expression, concomitant with the tumor reduction. Given the transient half-life of RNPs (∼24 h), it is not surprising that tumors treated with the dCas9-KRAB RNPs have a limited therapeutic window of approximately two weeks or of tumor inhibition phase (with ∼50% tumor reduction relative to no gRNA, vehicle control, or the same RNP formulations in presence of non-specific gRNA). After this interval, the dCas9-KRAB *EWSR1-FLI1* gRNA edited tumors showed no significant difference in tumor volume relative to controls. Thus, subsequent treatment cycles would be necessary to maintain the therapeutic effect, ideally in combination with other agents, as described above. A striking feature of our formulations is an extremely quick peak of action post-injection, which could facilitate a rapid therapeutic effect prior /and/ or in combination with chemotherapy or other treatment modalities. Lastly, we found that the dCas9-KRAB RNP previously characterized in cell lines was similarly able to reduce the growth of the ES-4 EWS patient-derived xenograft (PDX) by at least 50%. PDXs recapitulate the genetic and epigenetic landscape of human tumors, and they have been extensively and effectively utilized in several malignancies, including _EWS.91,137_

Recently, polymeric nanocarriers have gained more attention for protein, nucleic acids, and anticancer drug delivery, as evidenced by the growing number of clinical trials that have approached approval stages or already been approved by the FDA.^138–143^ However, despite these trials, polymeric nanoparticles have not yet been utilized for CRISPR delivery in humans. Here, we provide, for the first time, preclinical evidence for the delivery of dCas9-KRAB RNPs payloads using non-viral polymeric delivery systems for future clinical translation. Our study findings provide evidence for the successful delivery of highly customizable RNPs which could be tailored to a variety of oncogenic drivers to treat multiple cancers with unmet clinical needs.

Future work should also consider the targeting the formulations with specific targeted antibodies or ligands recognizing these tumors, such as anti-EGFR (as recently published with a lipid nanoparticle study targeting head-and neck squamous cell carcinoma^144^. In the case of EWS, potential targeted markers are CD99,^145^ GD2,^146^ LINGO1,^147^ and CD133^148^. These targeted approaches could potentially improve the homing and activity of the polymers *in vivo* and enable, ideally, the systemic administration of the RNPs ^139^. In the case of the targeting to the liver, a recent report showed successful intravenous delivery of lipid-RNP complexes for hepatic gene editing and T cell engineering ^42^. Future challenge will involve the intravenous delivery of RNP silencers for epi/genetic editing of solid tumors.

In conclusion, our work demonstrated a specific and customizable silencing approach using the CRISPR/dCas9-KRAB repression system that overcomes challenges associated with the delivery of CRISPR dCas9-KRAB, opening the door for precision oncology medicine and future clinical studies.

## Materials and Methods

Please refer to the supporting information document.

## Conflict of Interests

The authors declare no conflict of interest.

## Acknowledgments

This work was supported by a Cancer Council of Western Australia (CCWA) project grant and an Australian Excellence Award to PB. ST would like to acknowledge the financial support of the Australian Government Research Training Program (RTP), Sock it to Sarcoma, and CCWA PhD Top Up Scholarships. C.W.E. acknowledges funding by the State Government of Western Australia. The authors acknowledge Professor Vincent Rotello for providing mCherry-E_20_ protein. The authors would also like to acknowledge the technical assistance of the Australian Microscopy & Microanalysis Research Facility at the Centre for Microscopy, Characterisation & Analysis, The University of Western Australia, a facility funded by the University, State and Commonwealth Governments, and the Perkins Cancer Biobank.

## Author Contributions

Conceptualization: P.B. Methodology: P.B., S.T., C.W. (Christopher Wallis), L.D. (Larissa Daymond), C.W.E. (Cameron W Evans), L.W. (Louise Winteringham), K.S.I. (K Swaminathan Iyer), E.W. (Eleanor Woodward), P.H. (Peter Houghton), E.W. (Edina Wang), and M.N. (Marck Norret). Formal Analysis: S.T., and P.B. RNA-seq Analysis: S.T., and A.W. Investigation: P.B., S.T., C.W. (Christopher Wallis), C.W.E., E.W. (Eleanor Woodward), L.D., E.W. (Edina Wang), L.W. and S.G. Writing Original draft: S.T. Review and Editing: P.B., C.W (Charlene Waryah), A.T (Ash Tie), S.T., C.W.E., K.S.I., L.W., L.D., E.W. (Eleanor Woodward), P.H., C.W. (Christopher Wallis), M.N., and S.G. Figure collation and editing: Figure 1: S.T. and C.W. Figure 2: S.T. Figure 3: S.T. and A.W. Figure 4: S.T. Figure 5: S.T. Figure 6: S.T. and E.W. (Edina Wang). Figure 7: S.T. and L.D. Figure S1, Supporting Information 1: S.T. Figure S2, Supporting Information 1: S.T. Figure S3, Supporting Information 1: S.T. Figure S4, Supporting Information 1: S.T. and E.W. (Eleanor Woodward). Figure S5, Supporting Information 1: Ash Tie. Figure S6, Supporting Information 1: E.W. (Eleanor Woodward). Figure S7, Supporting Information 1: S.T. Figure S8, Supporting Information 1: S.T. Figure S9, Supporting Information 1: S.T. and C.W. Figure S10, Supporting Information 1: S.T. Figure S11, Supporting Information 1: S.T. and E.W. (Eleanor Woodward). Figure S12, Supporting Information 1: S.T. Figure S1, Supporting Information 2: S.T. Table S1, Supporting Information 1: S.T. and C.W. (Christopher Wallis). Table S2, Supporting Information 1: S.T. Supervision: P.B. Funding Acquisition for this projection: P.B. Project administration: P.B.

## Ethics Approval

All animal experiments were approved by the Animal Ethics Committee of the University of Western Australia (RA3/100/1718) and carried out in accordance with the Code of Practice for the Care and Use of Animals. Expansion of EWS PDX models and dCas9-KRAB RNP PDX experiments were approved by the Animal Ethics Committee at Harry Perkins Institute of Medical Research (AE253 and AE256) and conducted in accordance with the Australian Code for the Care and Use of Animals for Scientific Purposes.

## Data availability

The RNA-seq data have been deposited in the Gene Expression Omnibus public database under accession number GSE236399.

## Notes

### Competing Interest Statement

The authors have declared no competing interest.

