## Supplemental_Info for "Targeted epigenetic repression of oncogenic transcription factors via CRISPR/dCas9 locus-specific silencing"

### Supporting Information

**Table S1. gRNA sequences for *EWSR-FLI1* suppression.** Candidate gRNA sequences were identified using Benchling software and selected depending on specificity score > 70 and efficiency score >50. The start and end sites of each gRNA were specified based on the Genome Reference Consortium Human Build 38 (GRCh38). The start position of each gRNA was numbered according to the transcription start site of EWSR-FLI1 mRNA transcript variant 2 NM_005243.4 in GRCh38. PAM: protospacer-adjacent motif of spCas9 (NGG); F: forward strand guide; R: reverse strand guide.

| gRNA | Target Sequence | PAM | Strand direction R/F | Start | End | Specificity Score | Efficiency Score | Position |
|---|---|---|---|---|---|---|---|---|
| 1 | TGACTAATCCG CGGGCCGCG | GGG | R | 29267903 | 29267922 | 93.6 | 63.3 | -345 |
| 2 | AGGGACGGTC GCTGAAGAGT | CGG | R | 29267989 | 29268008 | 81.1 | 63.3 | -259 |
| 3 | CGCGAAGAAC GAGTAAGCGG | TGG | F | 29268173 | 29268192 | 94.6 | 64 | -95 |
| 4 | TGGCCCGATG GTCGCGCCCG | GGG | F | 29268226 | 29268245 | 96.2 | 57.5 | -42 |
| 5 | AGTTCCACCA TACTCACCCG | TGG | R | 29268347 | 29268366 | 87.8 | 74.4 | +99 |
| 6 | TAGCCGGAAC GCCCAAACTG | GGG | F | 29268383 | 29268402 | 86.9 | 64.7 | +115 |

**Table S2. Detection of potential matching sequences of *EWSR1-FLI1* gRNAs using Cas-OFFinder software.** Mismatches to target gRNA sequences are displayed in lowercase, bold, and underlined in the candidate off-target sequences. Location indicates the chromosomal position in GRCh38.

| gRNA | Candidate off-target sequences | Chromosome | Location | Direction | Proximity to regulatory regions | Potential gene affected |
|---|---|---|---|---|---|---|
| 1 | TG**g**CT**g**AT**t**CGCGG GCCGCGAGG | chr8 | 143840563 | + | 0.4 kb downstream of NRBP2 TSS, 25 kb upstream of SCRIB TSS, 11 kb upstream of PUF60 TSS, 32 kb downstream of EPPK1 TSS; within CpG island; in DNase I hypersensitive region; in H3K27ac, H3K4me3, H3K4me1 enriched regions | NRBP2<br>PUF60<br>SCRIB<br>EPPK1 |
| | TG**t**CTAA**c**CCGCGG GCCGC**a**TGG | chr3 | 9395253 | - | 32 kb downstream of THUMPD3 TSS; 2 kb upstream of SETD5 TSS; in DNase I hypersensitive region; in H3K4me3, H3K4me1 enriched regions | THUMPD3<br>SETD5 |
| 2 | AG**a**GACGG**g**CGC**g** GAAGAGTAGG | chr7 | 42026372 | + | None | None |
| | AGGGA**t**GGT**a**G**g**TG AAGAGTTGG | chr20 | 40639416 | - | 49 kb downstream of MAFB TSS; in DNase I hypersensitive region; in H3K4me3, H3K4me1 enriched regions | MAFB |
| | AGGGA**t**GGT**g**GCTG **t**AGAGTGGG | chr6 | 42521240 | + | 70 kb upstream of TRERF1 TSS, 43 kb upstream of UBR2 TSS; in DNase I hypersensitive region; in H3K4me3, H3K4me1 enriched regions | UBR2<br>TRERF1 |

| | | | | | | |
|---|---|---|---|---|---|---|
| 2 | AGGGAC**ta**TC**a**CTG AAGAGTTGG | chr3 | 95215722 | + | None | None |
| | **c**GGGA**t**GGT**a**GCTG AAGAGTTGG | chr3 | 129982011 | - | 7 kb downstream of TRH TSS; in DNase I hypersensitive region; in H3K4me3, H3K4me1 enriched regions | TRH |
| 3 | CG**g**GAAGAA**g**GAGT AAG**a**GGTGG | chr1 | 20654631 | + | 7 kb downstream of DDOST TSS; 21 kb downstream of PINK1 TSS; 63 kb downstream of KIF17 TSS; in DNase I hypersensitive region; in H3K4me3, H3K4me1 enriched regions | DDOST<br>PINK1<br>KIF17 |
| 5 | **t**GTTCCACC**g**TACTC ACCC**t**GGG | chr16 | 3428851 | - | 28 kb upstream of ZSCAN32 TSS; 28 kb downstream of ZNF174 TSS; 15 kb upstream of NAA60 TSS; 15 kb downstream of ZNF597 TSS; in DNase I hypersensitive region; in H3K4me3, H3K4me1 enriched regions | ZNF597<br>NAA60<br>ZSCAN32<br>ZNF174 |

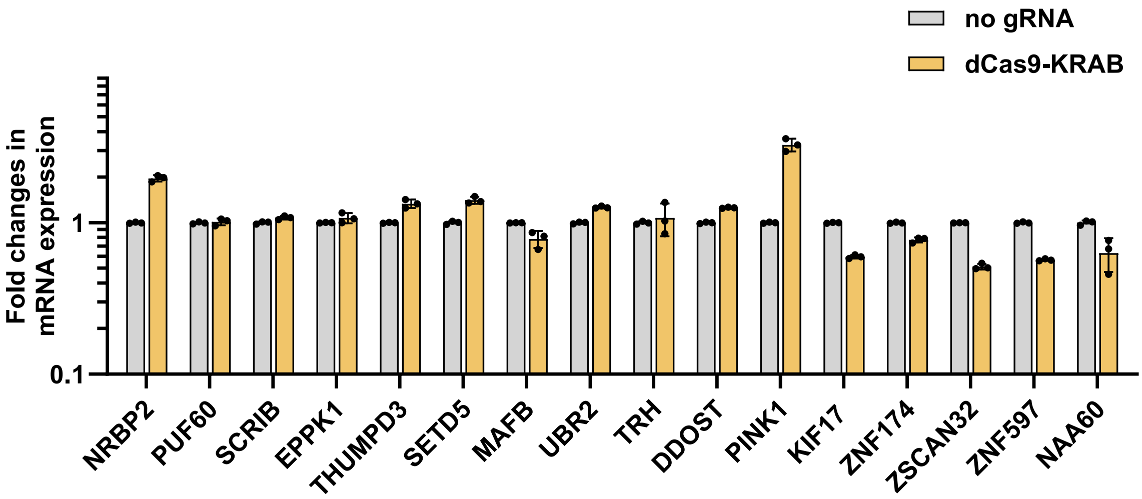


**Figure S1. The dCas9-KRAB system does not alter the regulation of predicted off-target genes.** qRT-PCR was performed to assess changes in the expression levels of mRNA of the top 16 predicted off-target genes in A673 cells expressing dCas9-KRAB delivered with gRNA mix compared to dCas9-KRAB with no gRNA. Data represent fold change in mRNA expression levels relative to dCas9-KRAB with no gRNA. No significant variation in expression was detected for all potential off-target genes. Ordinary one-way ANOVA with Dunnett multiple comparison test was applied to carry out statistical analysis. Statistical significance is assessed for biological triplicates and data are illustrated as mean values ± SD.

A)

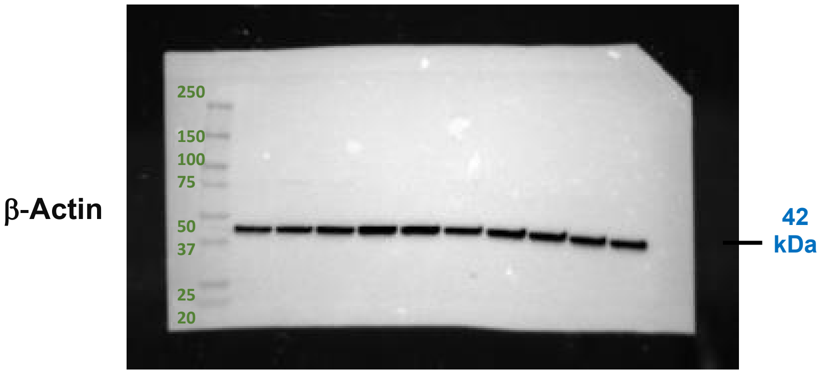
β-Actin
250
150
100
75
50
37
25
20
42 kDa

B)

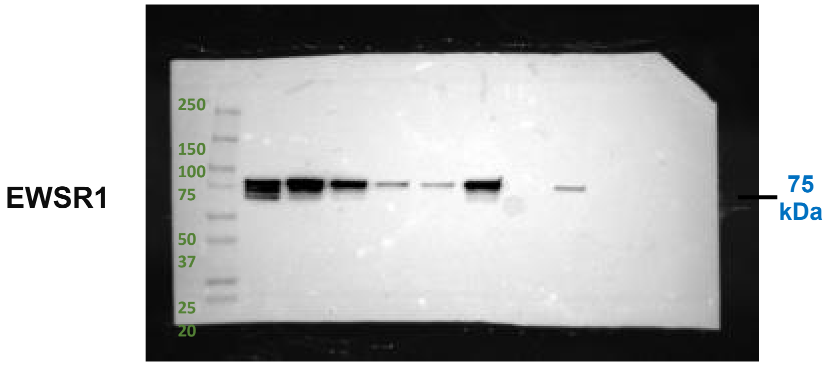
EWSR1
250
150
100
75
50
37
25
20
75 kDa

C)

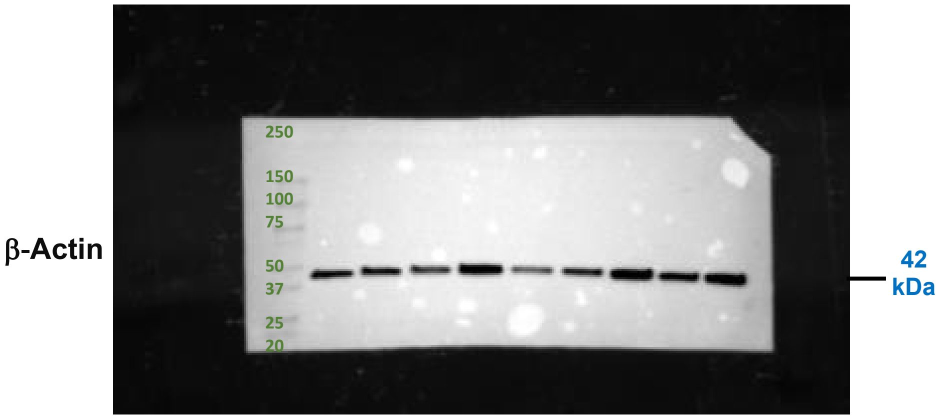
β-Actin
250
150
100
75
50
37
25
20
42 kDa

D)

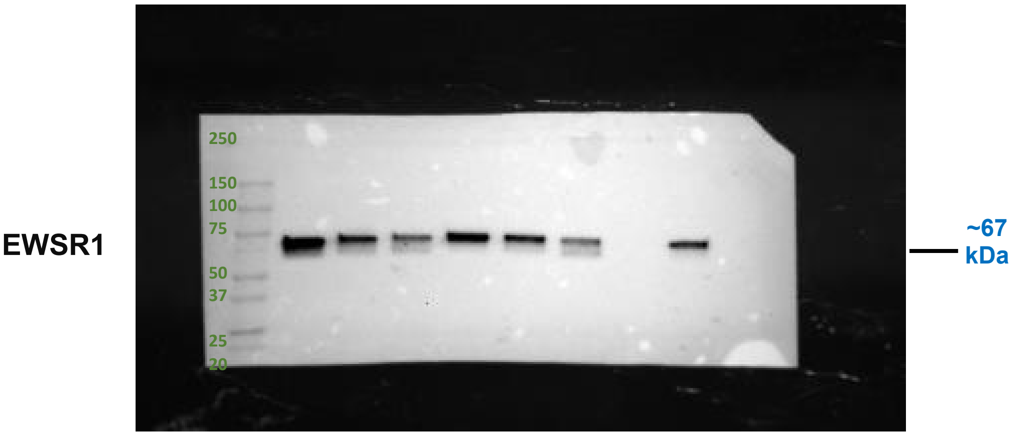
EWSR1
250
150
100
75
50
37
25
20
~67 kDa

**Figure S2. Original uncropped blots.** Uncropped full-length images of Western blot membranes correspond to (Figure. 1E-F) for b-Actin and EWSR1 protein expression in A673 (A,B) and RD-ES (C,D)**.** The size (kDa) of protein standards is displayed in green. The size (kDa) of EWS-FLI1 and b-Actin is presented in blue.

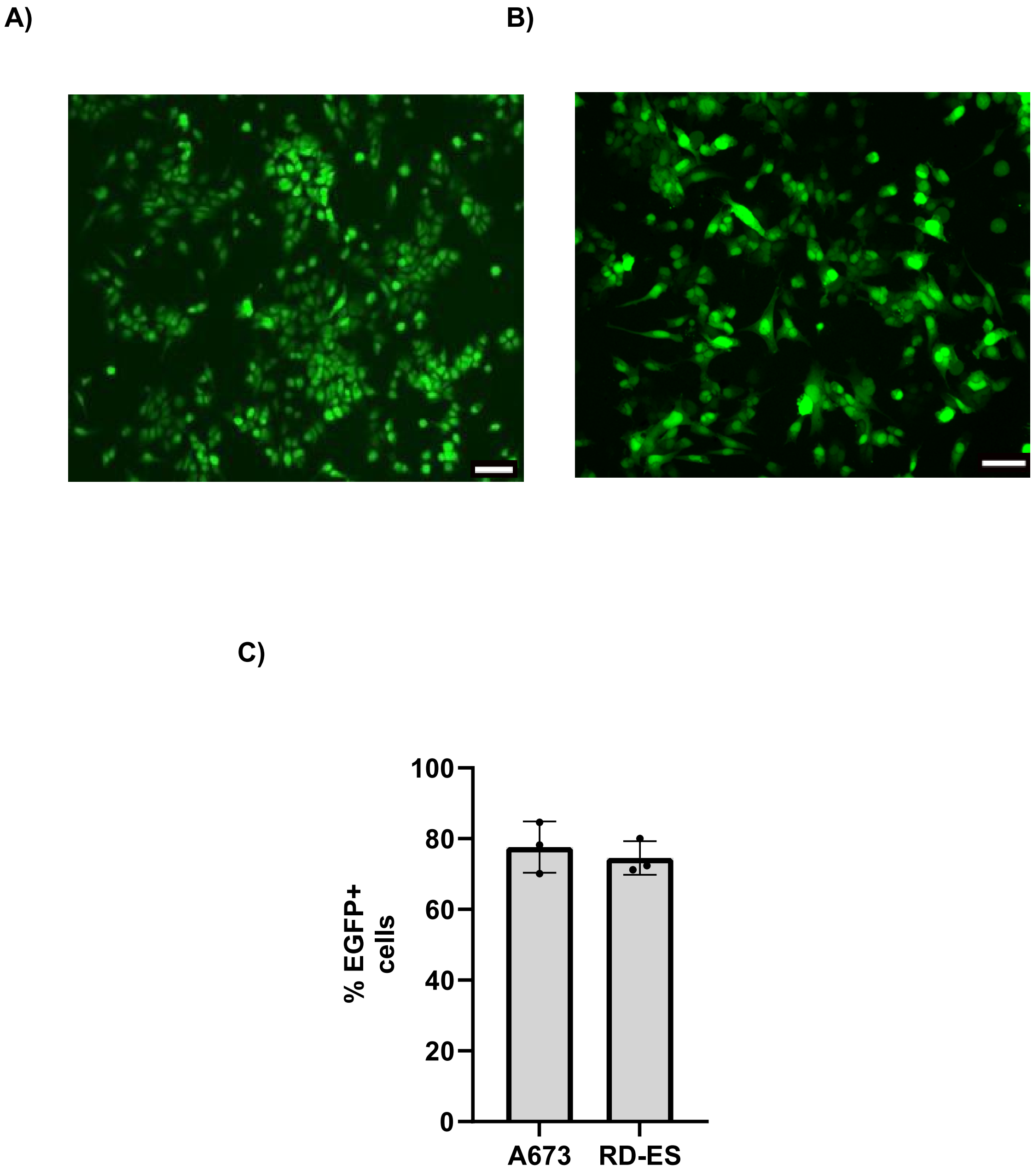


**Figure S3. Evaluation of lentiviral transfection efficiency by detecting EGFP expression levels in Ewing sarcoma cell lines.** A673 (A) and RD-ES (B) express high levels of EGFP (> 70% of cells) after lentiviral transfection with pCMV-EGFP plasmid (Addgene plasmid #11153) indicating highly efficient delivery of lentiviral transfection method (C). EGFP = green fluorescent protein. Scale bar = 100 µm.

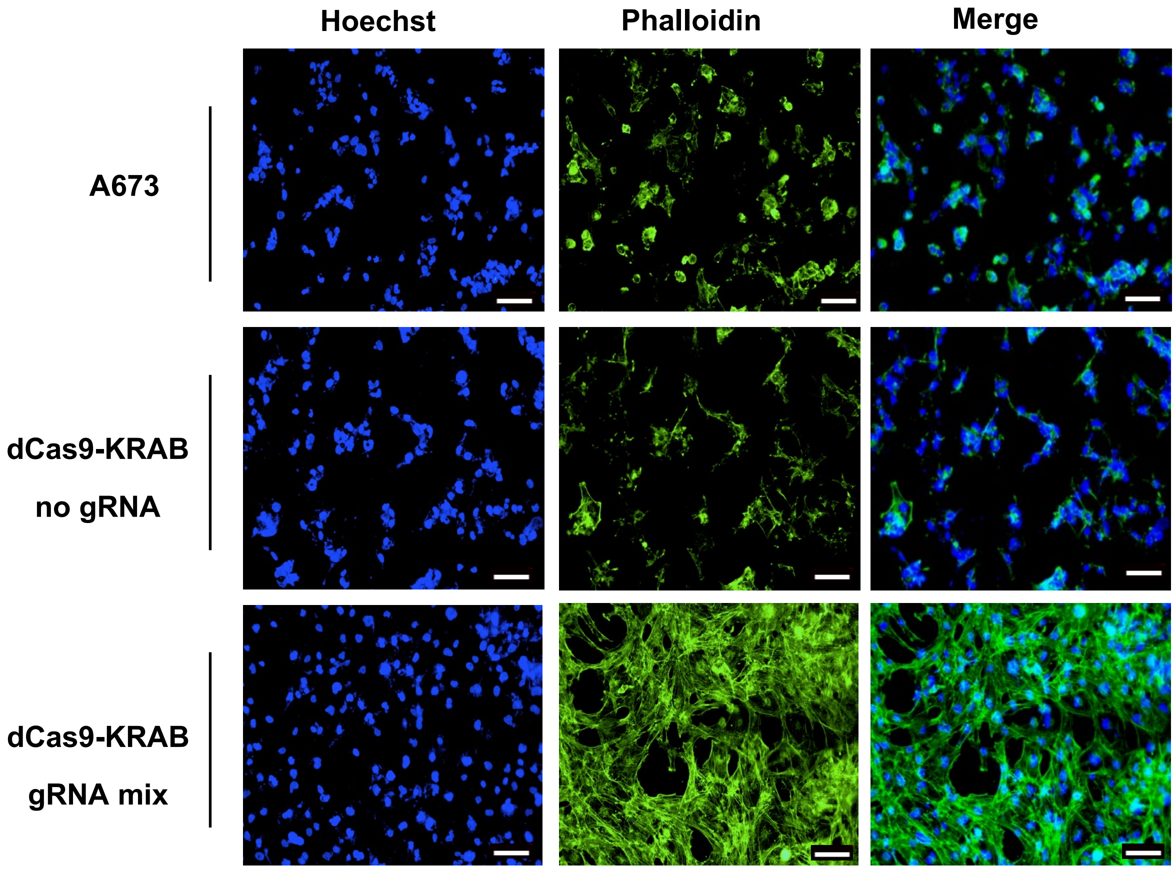


**Figure S4. Actin cytoskeleton organization post-EWS/FLI knockdown.** Representative images of Phalloidin-stained actin cytoskeleton and Hoechst staining in A673 non-transduced cells, transduced with dCas9-KRAB-no gRNA or with dCas9-KRAB-gRNA mix cells. Scale bar represents 100 µm.

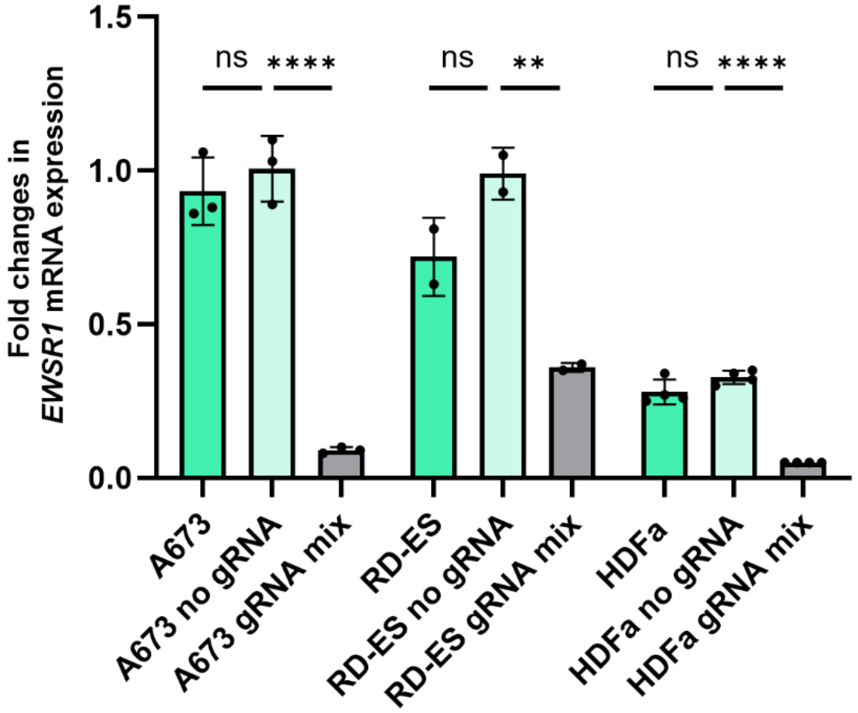


**Figure S5:** Endogenous *EWSR1* expression in Ewing sarcoma (A673, RD-ES) and normal primary human dermal fibroblast (HDFa) cell lines. Lentiviral delivery of dCas9-KRAB with either no gRNA or gRNA mix was performed across the cell types. In Ewing sarcoma cell lines, both *EWSR1* and *EWSR1-FLI1* fusion transcript are detected. The wildtype HDFa cells display low endogenous *EWSR1* expression. All values are normalized to A673 no gRNA. Data represent mean ± SD from n=2-4 biologically independent experiments. Statistical significance was determined using Student's *t*-test relative to A673 no gRNA. **$p < 0.01$; ****$p < 0.0001$.

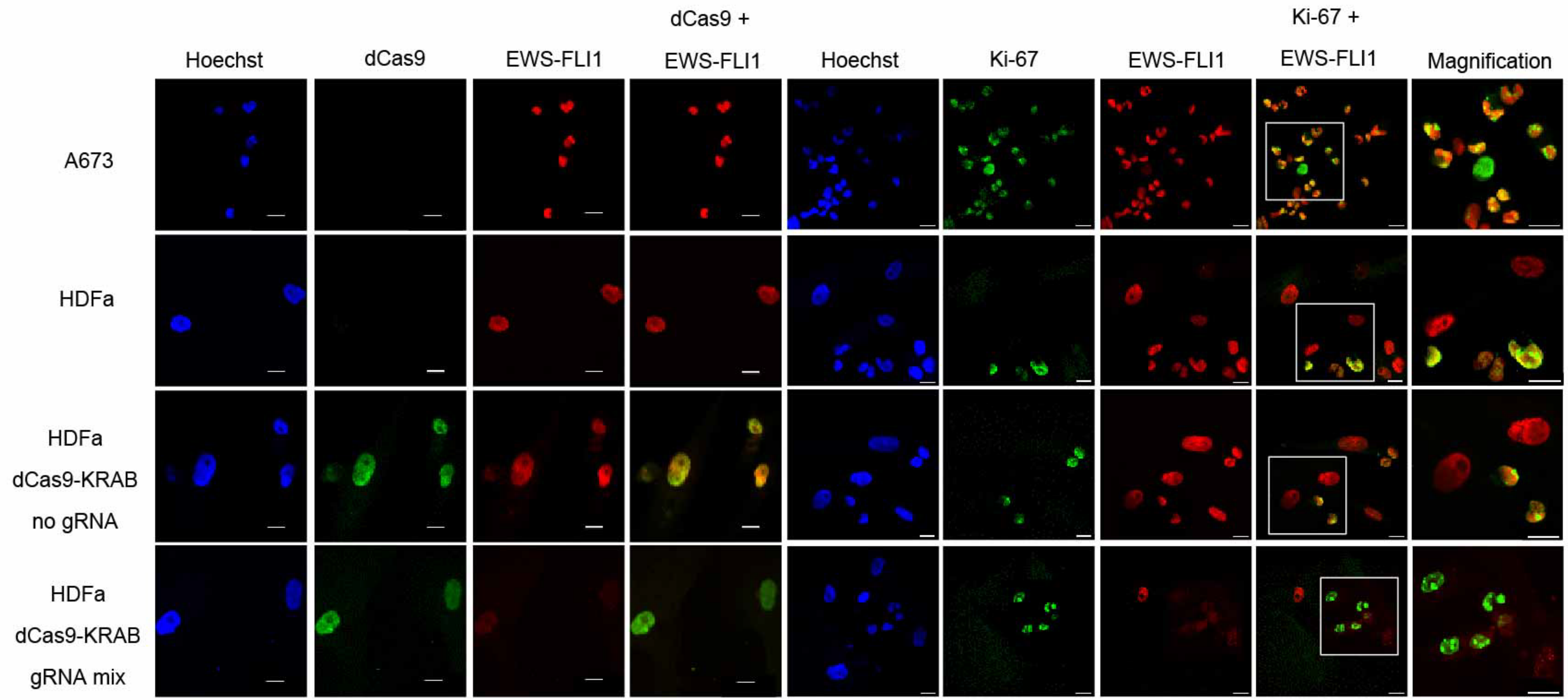


**Figure S6. CRISPR/dCas9-KRAB construct did not affect proliferation in HDFa cells.** Representative immunofluorescence staining of Ki-67 (green), dCas9 (green), EWSR1 (red) and nuclei (Hoechst 33258, blue) in A673 cells, HDFa cell and HDFa transduced with CRISPR constructs. Scale bar represents 20 µm.

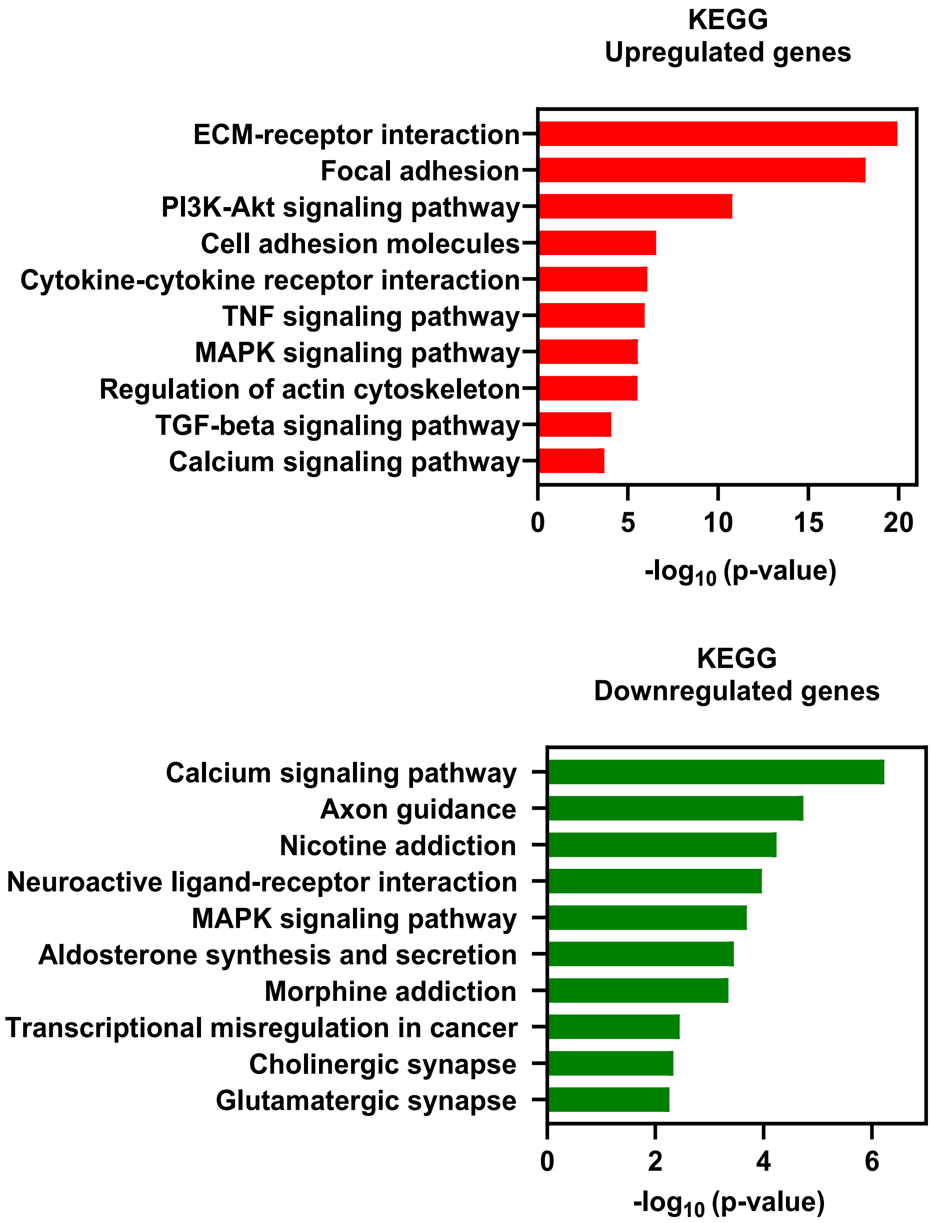


**Figure S7. Visualization of top enriched KEGG pathways.** The top significant pathways with $p < 0.01$ are presented. The *y*-axis represents the pathway, and the *x*-axis represents -$\log_{10}$ (p-value) of enrichment.

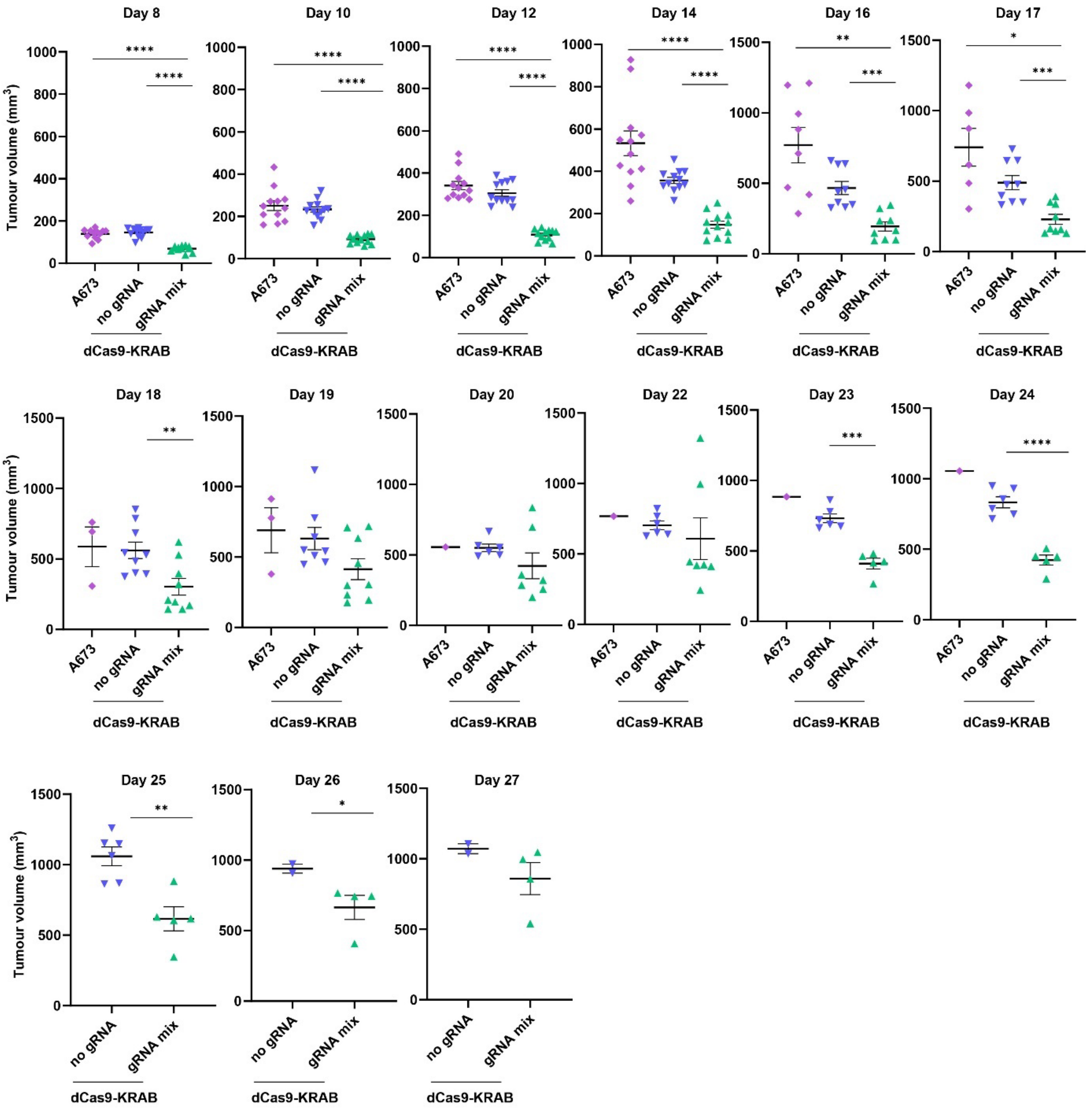


**Figure S8. Suppression of *EWSR1-FLI1* in Ewing sarcoma xenograft models using dCas9-KRAB with gRNA mix.** Scatter plots demonstrating the tumor volume ($mm^3$) of an individual mouse from day 8 to day 27 post-implantation. Data are plotted as mean values ± SEM, and *p*-values were determined using unpaired *t*-test (*$p < 0.05$, **$p < 0.01$, ***$p < 0.001$, ****$p < 0.0001$).

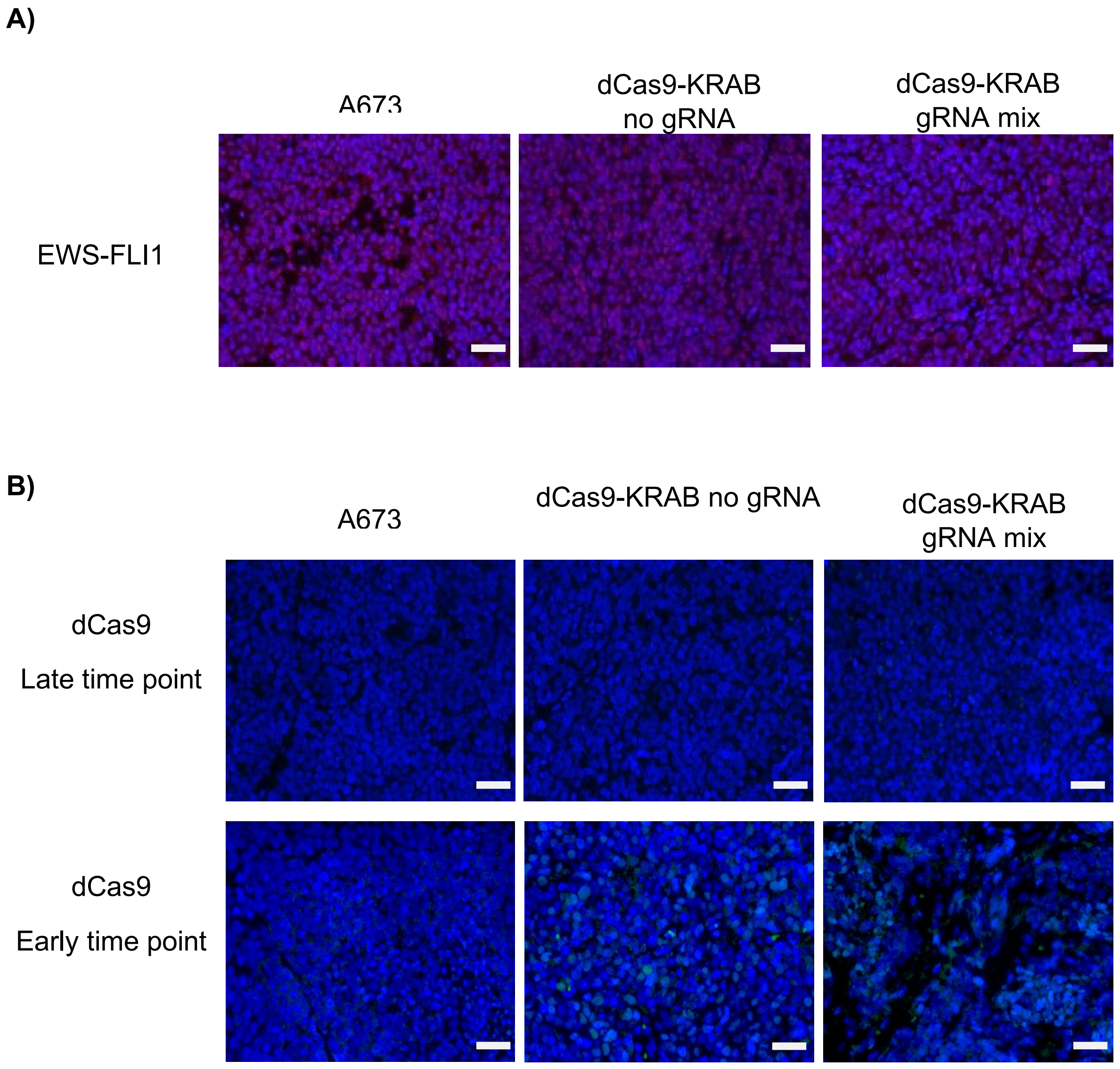


**Figure S9. Suppression of *EWSR1-FLI1* in EWS A673 transduced xenograft models by dCas9-KRAB.** Representative sections of tumors collected from (n = 3 mice per group) when tumors reached 1000 $mm^3$ stained with EWS-FLI1 (red) (A) and dCas9 (green) (B) antibodies (samples collected at early time point were used as positive control for dCas9 staining). Scale bars =50 μm.

A)

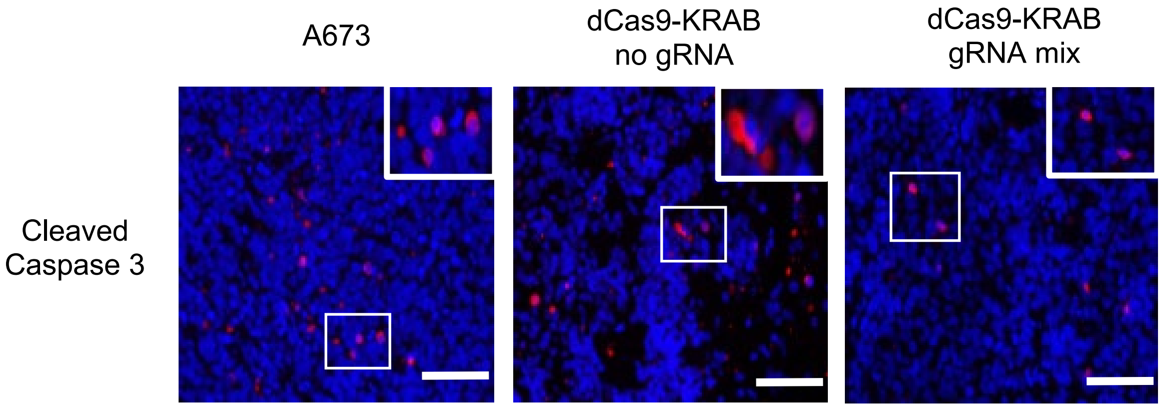


B)

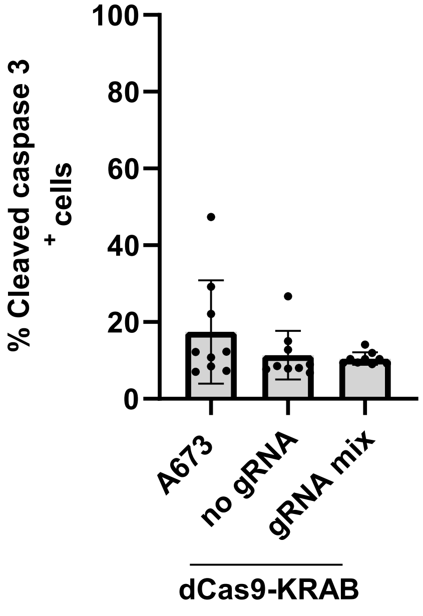


**Figure S10.** A) Representative sections of tumors collected at day 15 from (n = 3 mice per group) stained with apoptosis marker (anti-cleaved caspase 3) (red). B) Quantification of this marker. Scale bars represent 50 μm. Data are plotted as mean values ± SD, and statistical analysis was carried out using multiple unpaired *t*-tests.

A)

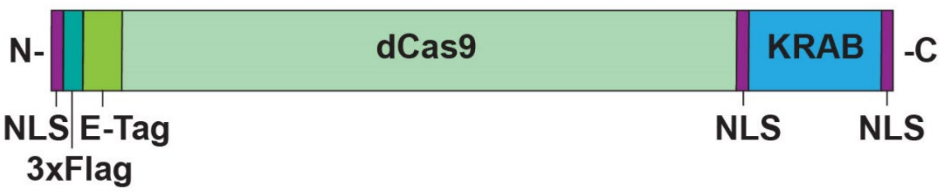


B)

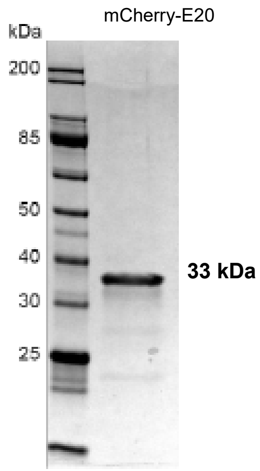


C)

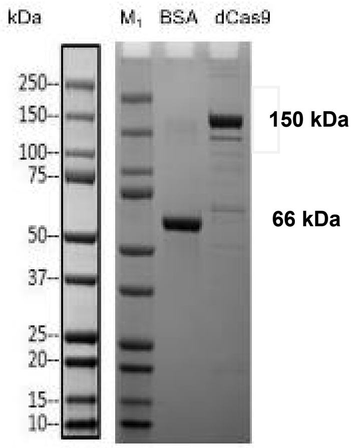


**Figure S11.** A) Schematic representation of the dCas9-KRAB protein. B,C) 10% SDS-PAGE gel reflecting the purity of the dCas9 and mCherry-$E_{20}$ proteins. Both proteins were harvested at high purity (nearly 75%) using a HisPur cobalt column. B) A strong band for dCas9 protein was observed at ~ 150 kDa. The BSA protein (control) band was observed at ~ 66 kDa. Lane $M_1$: Protein Marker (carried out by GenScript). C) A strong band for mCherry-$E_{20}$ protein was detected at ~ 33 kDa containing the mCherry protein with 6× His tags and 20× E-tags (adapted from Kretzmann *et al.*[1]).

A)

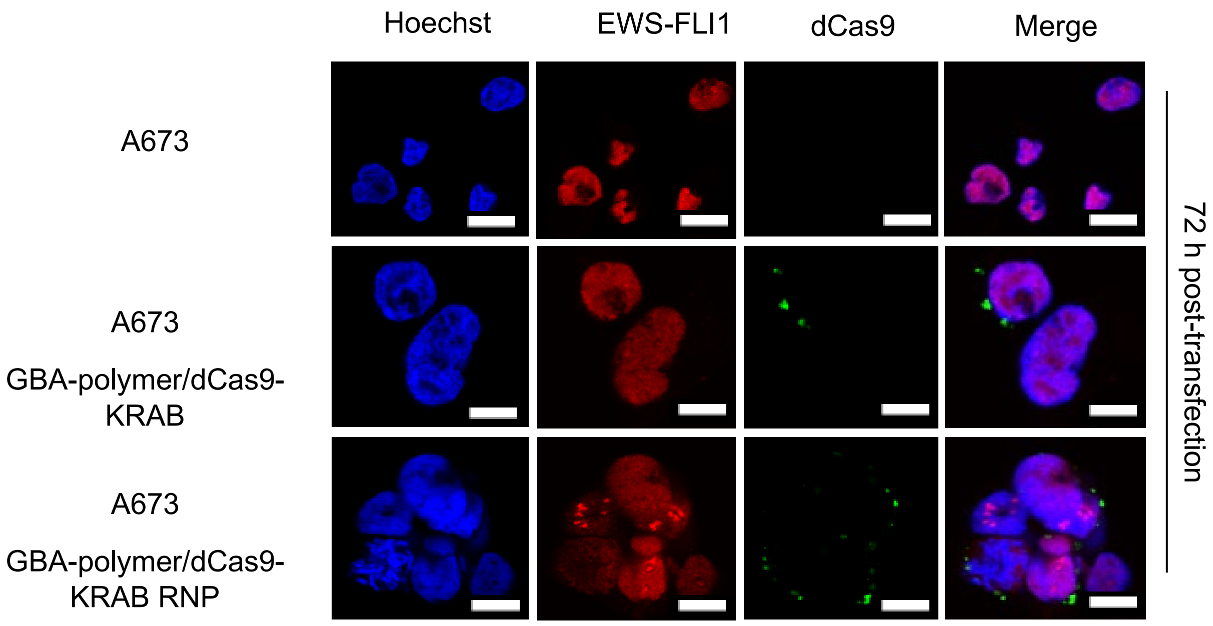


B)

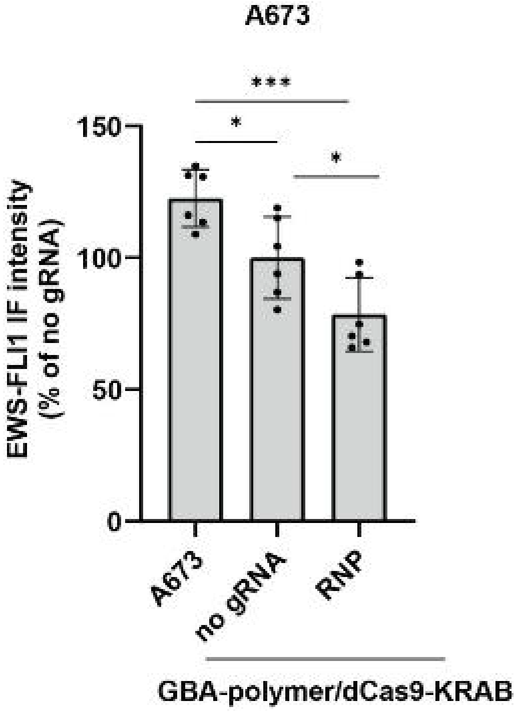


**Figure S12. Effects of GBA-polymers/dCas9-KRAB in suppressing EWS-FLI1 72h post-transfection in A673.** A) Representative images for dCas9-KRAB protein delivery to A673 cells using the PAMAM- polymeric nanocarrier at mass ratio 1.5:1 (polymer: protein) after 72

h of transfection. EWS-FLI1 immunostaining (red), dCas9 immunostaining (green) and nuclear Hoechst staining (blue). Scale bar represents 10 µm. B) Mean fluorescent intensity (MFI) of EWS-FLI1 protein after 72 h of transfection by polymer. Data are normalized to the dCas9-KRAB with no gRNA and presented as mean values ± SD. *p*-values are determined by multiple unpaired *t*-test (***$p < 0.001$, *$p < 0.05$). n = 3 biologically independent experiments.

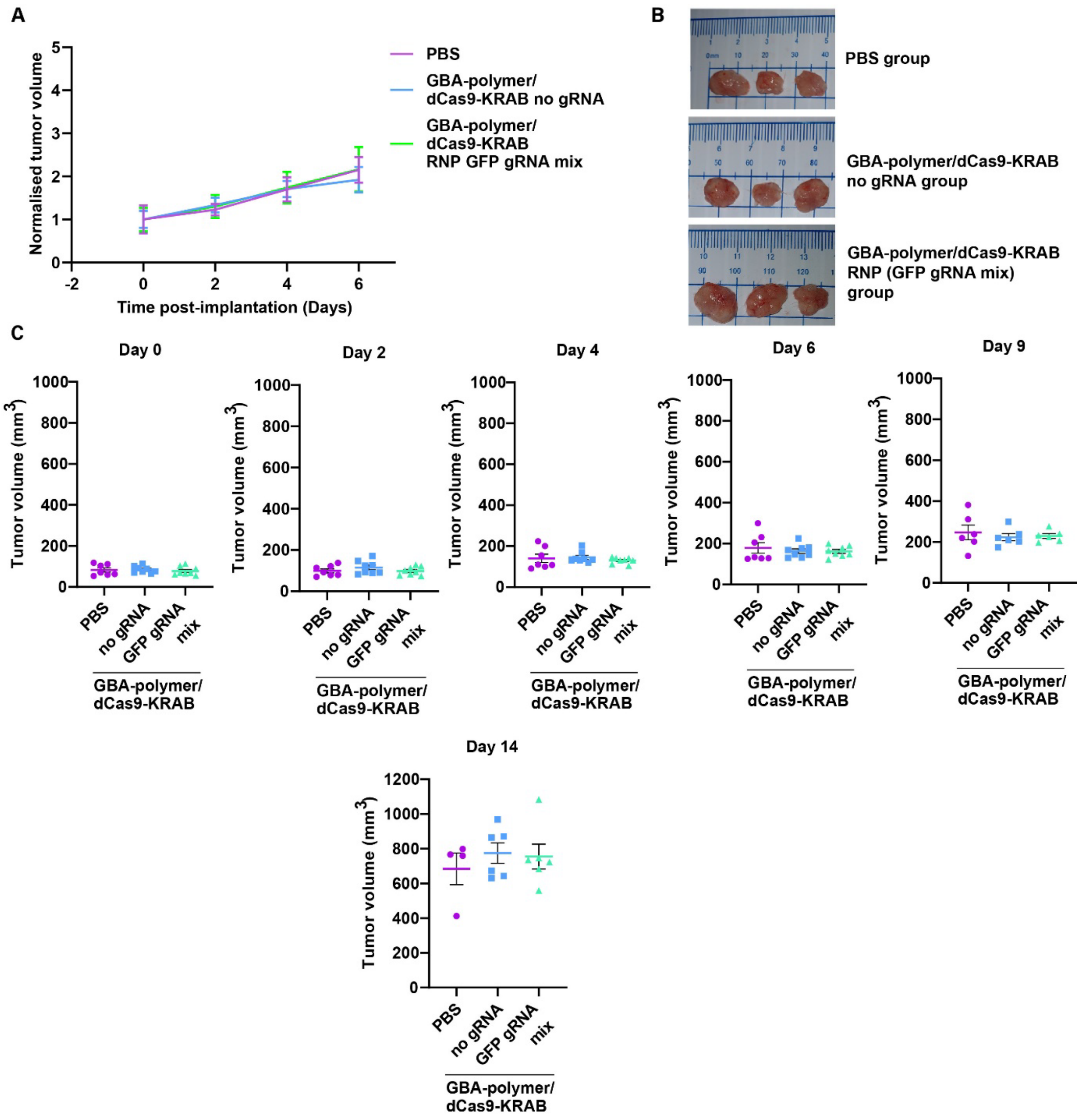


**Figure S13.** ***In vivo*** **tumor growth experiment using A673 xenograft mouse models. A)** Normalized mean tumor volumes in NSG mice treated intratumorally with either GBA-polymer/dCas9-KRAB RNP (GFP gRNA mix) with GFP gRNA (n = 8, green), GBA-polymer/dCas9-KRAB with no GFP gRNA (n = 8, blue), or vehicle (PBS) (n = 7, purple). Mice received four doses in total, one injection every 48 h. Tumor volumes were normalized to day 0 post-injection and presented as mean values ± SEM. B) Images of representative of ex vivo A673 tumors extracted at day 15 post-injection. C) Corresponding box plots of tumor volume measurements at days 0, 2, 4, 6, 9 and 14 post-injection. Statistical significance is calculated using multiple unpaired *t*-test.

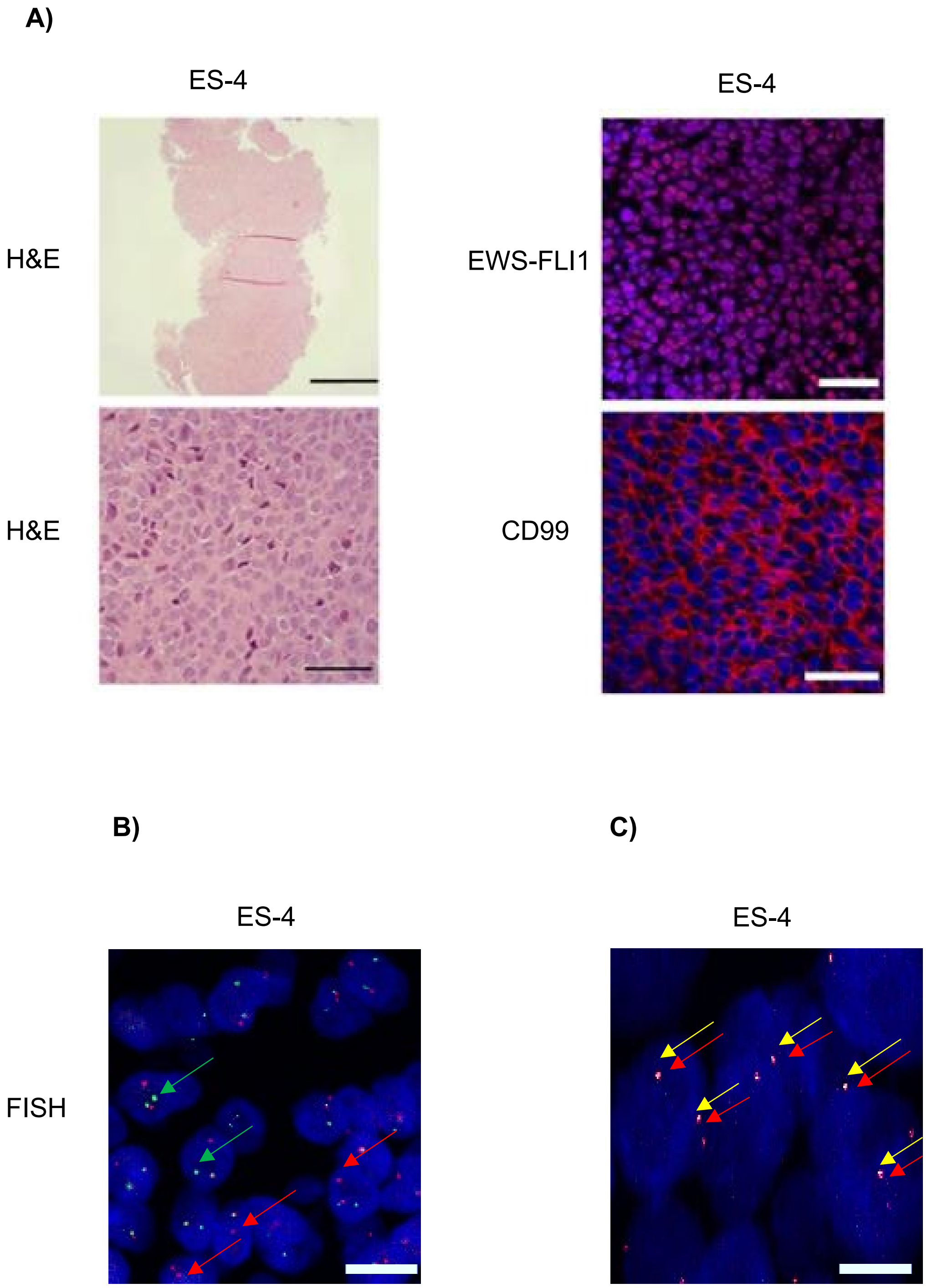
A)
ES-4
ES-4
H&E
EWS-FLI1
H&E
CD99
B)
C)
ES-4
ES-4
FISH

**Figure S14. Histologic and immunohistochemical features of ES-4 PDX tumor. A)** Immunohistochemistry analysis of PDX tumor biopsies stained with Hematoxylin and eosin, anti-EWSR1, anti-CD99, and Hoechst. Images are at 4×, 20× and 40× magnification. B) Dual-colour fluorescence in situ hybridization utilizing break-apart *EWSR1* probes. Separated red and green signals in nuclei of EWS cells denote a rearrangement of the *EWSR1* gene at 22q12. A fused yellow signal represents the site of the *EWSR1* gene. C) Dual-colour fluorescence in situ hybridization using *EWSR1* and *FLI1* probes. Separated red and yellow signals in nuclei of EWS cells denote the *EWSR1* and *FLI1* genes. A fused orange signal represents the site of the *EWSR1-FLI1* rearrangement. Images were obtained by a confocal microscope at 60× magnification. Scale bar represents 10 μm.

### Materials and Methods

#### *Guide design and cloning*

Repression of the *EWSR1-FLI1* fusion gene was achieved using a CRISPR/dCas9 fused to the KRAB effector domain. The plasmid pLV-hU6-sgRNA-hUbC-dCAS9-KRAB-T2A-Puro (a gift from Charles Gersbach, Addgene, #71236), is a third-generation lentiviral vector that contains a single-guide RNA (sgRNA) driven by a U6 promoter and 3X FLAG tagged dCas9 fused to the KRAB at the C-terminus. From hereafter, this plasmid is referred to as pLV-dCas9-KRAB.

A panel of six sgRNA with optimized on-target and minimized off-target scores (Benchling) were identified within a 600bp region spanning 400 bp upstream and 200 bp downstream of the transcription start site of the *EWSR1* gene. The corresponding guide oligos were synthesized by Integrated DNA Technologies (IDT) and cloned into the pLV-dCas9-KRAB vector. Successful insertion of sgRNA into the vector was verified by restriction enzyme digestion and Sanger Sequencing (ARGF, Perth). Unless otherwise stated, all sgRNA experimental conditions contained an equal total plasmid concentration of the six sgRNA. As a negative control, a non-targeting pLV-dCas9-KRAB plasmid lacking sgRNA was used.

#### *Cell lines*

The human embryonic kidney 293T (HEK293T, CRL-3216), primary human dermal fibroblast (PCS-201-012) and EWS cell lines (A673; CRL1598 and RD-ES; HTB-116) were obtained from American Type Culture Collection (ATCC). A673 and HEK293T cells were maintained in Dulbecco's Modified Eagle's Medium (DMEM; high glucose with pyruvate) supplemented with 10% heat-inactivated fetal bovine serum (HI-FBS; Gibco) and 1% Antibiotic-Antimycotic (Anti-Anti; Gibco). RD-ES cells were maintained in Roswell Park Memorial Institute (RPMI-1640) medium, supplemented with 10% HI-FBS and 1% Anti-Anti. The cells were maintained at 37°C and 5% $CO_2$ in a tissue culture incubator.

#### *Lentiviral transduction*

All lentiviral experiments were performed in accordance with the approval from the Perkins Institutional Biosafety Committee (NLRD004/2017) and Office of Gene Technology Regulator (NLRD008/2020). Lentiviral particles were produced in HEK293T cells by co-transfecting with third generation packaging plasmid VSV-G (Addgene, #12259) and Gag/Pol (Addgene, #12260) with pLV-dCas9-KRAB as previously described. Four days following viral transduction, 1ug $ml^{-1}$ and 2ug $ml^{-1}$ of puromycin (Gibco) was used to select transduced A673 and RD-ES cells, respectively. After conducting off-target analysis, 3 gRNA were selected (referred to as "gRNA mix"). Cells transduced with the non-targeting vector (no sgRNA) are referred to as “no-gRNA” controls. Lentiviral transduction efficiency was assessed by a reporter lentivirus pLV-eGFP (Addgene, #36083).

*RNA extraction and RT-qPCR*

Total RNA was extracted using QIAzol Lysis Reagent (QIAGEN) following the manufacturer’s protocol. The purify and concentration of RNA was assessed using NanoDrop™ spectrophotometer (ThermoFisher Scientific) before 2 mg of total RNA was used for cDNA synthesis using the High-Capacity cDNA Reverse Transcription Kit (Applied Biosystems). Relative transcript expression levels were determined using TaqMan Probes (Applied Biosystems; Table S2) or custom primers with the QuantiFast SYBR Green PCR Kit (Thermo Fisher Scientific; Table S2). PCR was performed on the ViiA 7 Real-Time PCR System (Applied Biosystems). Relative quantification of gene expression was normalized to the housekeeping gene GAPDH, using comparative $2^{-\Delta\Delta CT}$ method.

*RNA-Sequencing and analysis*

Total RNA was extracted using QIAzol Lysis Reagent (QIAGEN) as per the manufacturer’s protocol. RNA sequencing libraries were prepared using TruSeq Stranded mRNA Library Prep Kit (Illumina, San Diego, USA) and sequenced to generate 50 bp paired-end reads on the Illumina HiSeq2500 platform (Genomic WA). Adapter sequences were trimmed via BBDuk and aligned by STAR to the human assembly GRCh38. Differential gene expression was performed using DESeq2 (Genomic WA). iDEP.96 webtool was utilized for hierarchical clustering analysis and heatmap generation of the most variable differentially expressed genes ($|\log_2 FC| > 1$, adjusted $p < 0.05$). Gene Set Enrichment Analysis (GSEA) was performed to identify significantly enriched biological pathway.

*Protein extraction and Western blot*

Protein was extracted from pelleted cells using Cell Lysis Buffer (Cell Signaling Technology) according to the manufacturer’s instruction. Cell lysates were sonicated for 10 s at 10 mA followed by centrifugation at 13,000 rpm for 10 min at 4°C. The supernatant was collected, and protein concentration was measured using DC™ Protein Assay (Bio-Rad), as per manufacturer’s protocol. Briefly, equal amounts of proteins were resolved on a 4 - 20% Mini-PROTEAN TGC Precast Gel (Bio-Rad) and transferred onto a PDVF membrane using the Trans-Blot Turbo Transfer System (Bio-Rad). Membranes were blocked with 5% (w/v) skim milk in TBST (20 mM Tris, 150mM NaCl, 0.1 Tween 20) and washed before being incubate overnight at 4°C with primary antibodies (Table S4) diluted in blocking buffer. After washing, membranes were incubated for 1 h at room temperature (RT) with secondary HRP-conjugated antibodies diluted in blocking buffer. Protein bands were visualized using enhanced chemiluminescence reagent (Millipore) and imaged using the iBright FL1500 Intelligent Imaging System (ThermoFisher Scientific). Images were processed using the iBright Analysis Software (ThermoFisher Scientific).

*Immunofluorescence staining of cultured cells*

Cells were seeded onto a poly-L-lysine coated coverslip and incubated overnight at 37°C and 5% CO2. Cells were then fixed with 4% paraformaldehyde (PFA) in PBS, washed with PBS then blocked with 5% Goat Serum/0.3% Triton X-100/PBS for 1 h at RT. Blocking solution was removed, and cells were incubated for overnight at 4°C with the appropriate primary antibody (Table S5) in 1% BSA/PBS. Cells were then incubated for 1 h at RT with secondary antibodies Alexa Fluor 488 goat anti-mouse (ThermoFisher Scientific), Alexa Fluor 594 goat anti-rabbit (ThermoFisher Scientific), and Hoechst (Sigma-Aldrich) in 1% BSA/PBS. Coverslips were mounted with SlowFade Diamond Antifade Mountant (Molecular Probes, Eugene, Oregon). Images were captured using a fluorescent microscope (Nikon) and processed using NIS-Elements Software.

*Cell viability assay*

CellTiter-Glo® 2.0 (Promega) was used to quantify cell viability. Briefly, A673 and RD-ES cells were seeded into a 96-well optical bottom white tissue culture plate at 2500 cells/well and 2600 cells/well respectively. The cells were cultured at 37°C and 5% $CO_2$ in a humidified tissue culture incubator. Cell viability was evaluated at 1 h, 2 h, 48 h, 72 h and 96 h post-seeding of the cells. The luminescent signals were recorded using EnVision 2102 Multilabel Reader (PerkinElmer).

*Soft agar colony forming assay*

The transduced A673 and RD-ES plus wildtype were evaluated for colony formation using the soft agar assay. Colonies were maintained in culture over 3 weeks with the culture media replaced every three days. Cells were stained with 0.5 mg $mL^{-1}$ MTT solution for 4 hours before imaging and quantification using microscope IX-71 (Olympus) and ImageJ software, respectively.

*Scratch wound closure migration assay*

Cells were cultured in IncuCyte® ImageLock 96-well plates (Essen Bioscience) until a monolayer was formed. The cells were serum starved for 24 h before a scratch on the cell monolayer was introduced using the Essen Bioscience 96-pin WoundMaker. All images and wound confluence measurements were performed on the IncuCyte® live-cell imager and processed with the IncuCyte® Zoom software (Essen Bioscience).

*Polymer nanoparticle synthesis and dendronization*

*Synthesis*: A statistical random polymer containing the monomers glycidyl methacrylate (GMA, 1) and hydroxyethyl methacrylate (HEMA, 2) was synthesized using atom transfer radical polymerization (ATRP) methodology. Briefly, inhibitors were removed from the monomers by passing through a basic alumina column. Subsequently, GMA (1.36mL, 10.0mmol) and HEMA (3.64mL, 30.0mmol) were taken up in MeOH (15mL), followed by addition of 2-(4-morpholino)ethyl 2-bromoisobutyrate initiator (Me-Br, 210μL, 1 mmol) and the solution degassed using 'freeze-pump-thaw' (3 cycles) method. Copper (I) bromide (CuBr, 76mg, 0.53 mmol) was added, followed by 2,2'-bipyridine (bpy, 210mg, 1.34 mmol) and the reaction heated to 80 °C under argon for 2 h. Reaction was opened to air and additional MeOH (15 mL) added. The solution was added dropwise to $Et_2O$ (300mL) and the product collected by filtration, before further purification by repeated precipitation by redissolving in MeOH (50ml), then addition to $Et_2O$ (300mL). After 3 precipitation cycles the product was dried overnight under vacuum to give Poly($HEMA_y$-ran-$GMA_x$) (3) (2.69g).

Copolymer (3) (1.0g) was heated to 60°C with sodium azide ($NaN_3$, 0.9g) and ammonium chloride ($NH_4Cl$, 750mg) in DMF(20mL) for 72h, then cooled to RT. Product obtained by filtration after dropwise addition of supernatant to $Et_2O$. Further purification by repeated precipitation by redissolving in MeOH (50ml), then addition to $Et_2O$ (300mL). After 3 precipitation cycles the product was dried overnight under vacuum to give Poly($HEMA_y$-ran-$Azido_x$) (4) (1.0g).

*Dendronization*: Generation 4.5 PAMAM dendron containing an alkyne focal point was synthesized. Dendron (702mg, 114 μmol) was dissolved in DMF (7mL) then added to 4 (33mg, 57μmol), followed by addition of pentamethyldiethylene triamine (PMDETA, 30μL, 144μmol). Solution degassed via 'freeze-pump-thaw' (3 cycles), then CuBr (25 mg, 174 μmol) added, and the reaction stirred under argon for 72h at RT. After completion the reaction mixture was transferred to dialysis tubing (10000MWCO) and dialysed against DI water (4x4L) for 24h, and the product collected by lyophilisation. The product was taken up in MeOH (2mL), then added dropwise to a cooled solution of ethylenediamine (2mL) in MeOH (2mL) and left stirring under argon for 5 days. After completion the reaction mixture was transferred to dialysis tubing (10000MWCO) and dialysed against DI water (4x4L) for 24h, and the product (5) collected by lyophilisation.

Partial guanidinylation of amino groups on dendronized polymer 5 was adapted from methods described by Chang *et al.*[156]. Briefly, guanidinobenzoic acid (GBA, 18.3mg, 84.6μmol ), 1-ethyl-3-(3-dimethylaminopropyl) carbodiimide (EDC, 9.8mg, 51.1μmol) and N-hydroxysuccinimide (NHS, 16.3mg, 14.2μmol) were dissolved in DMF (2mL) and stirred under argon at rt for 6 h. Separately, polymer (20mg, equivalent to 80μmol -$NH_2$) and triethylamine

(TEA, 50μL) was dissolved in PBS (pH = 7.4, 1mL) and added to the reaction mixture. Reaction left stirring at rt for 5 days under argon. After completion the reaction mixture was transferred to dialysis tubing (10000MWCO) and dialysed against DI water (4x4L) for 24h, and the product (6) collected by lyophilisation.

*Characterisation*: The average size (hydrodynamic diameter) and charge of polyplexes, dynamic light scattering (DLS) and zeta potential measurements were conducted using Zetasizer Nano ZS with He-Ne laser beam at a fixed scattering angle of 173°. Modified mCherry (mCherry-$E_{20}$) containing 20 glutamic acid tags was used a positive control when complexing with the polymer. Different GBA-polymer: protein mass ratios were prepared by diluting the polymer in PBS with a total of 40 ml and then mixing each sample with 2.5 mg of either mCherry-$E_{20}$ or dCas9-KRAB. Prior to measurements, the resulting polyplexes were incubated for 30 min at RT. All measurements were carried out in triplicate at 25°C with refractive index of 1.515 and 1.33 for PGMA and PBS respectively. Results were analysed using Malvern DTS 7.03 software and presented as mean ± standard deviation of the modal size peak. A mass ratio of 1.5 was identified to the ideal ratio for mCherry-$E_{20}$ and dCas9-KRAB as it has <500nm and a stable zeta potential (Figure S17).

*In vitro dCas9-KRAB delivery of polymeric nanoparticles*

gRNAs targeting *EWSR1-FLI1* and non-specific GFP gRNAs were ordered from IDT. A673 and RD-ES cells were seeded at a density of 2 × $10^5$ cell $mL^{-1}$ on poly-L-lysine coated glass coverslips in 24-well plates to achieve 80% confluency at the time of transfection. For GBA-polymer/mCherry-E20 nanoassembly preparation, GBA-polymer and mCherry-E20 were mixed at the optimized ratio (1.5: 1 mass ratio GBA-polymer to mCherry-E20) and incubated at RT for 30 min. For GBA-polymer/dCas9-KRAB RNP complex formation, dCas9-KRAB protein was diluted in Opti-MEM, mixed thoroughly with gRNAs solution at a mass ratio of 5:1 dCas9 to gRNAs, and incubated at RT for 30 min to form dCas9-KRAB RNP. After 30 min, the polymer was diluted in Opti-MEM, mixed with dCas9-KRAB RNP at a mass ratio of 1.5:1 and incubated at RT for 30 min. A GBA-polymer/dCas9-KRAB with no gRNA complex was also assembled using the same protocol, substituting the volume of gRNA with OptiMEM. Prior to nanoparticle delivery, cells were washed twice with PBS and replaced with 170 μL Opti-MEM. Polyplexes were added onto the cells and then incubated for 4 h, before being added with 280 μL of complete medium. Cells were fixed after 24 h for analysis of expression of mCherry or dCas9-KRAB protein through fluorescence microscopy.

*Animal experiments*

All animal experiments were performed in accordance with the protocols approved by the Animal Ethics Committee of the University of Western Australia (RA/3/100/1159 and

RA/3/100/1687) and the Australian code for the care and use of animals for scientific purposes. 4-week-old female NOD-scid-IL2Rgamma null (NSG) immunocompromised mice were purchased from Animal Resources Centre (Canning Vale, Australia). We performed power analysis using repeated-measures ANOVA with an F test for between-within subjects with Greenhouse-Geisser correction. The number of mice per group for each experiment provided sufficient power (>0.8) to detect an effect size of one. The mice were randomly assigned to different experimental groups.

*In vivo tumor growth assay of EWSR1-FLI1 silenced cell lines*

For the generation of the *EWSR1-FLI1* engineered xenografts, 1.5 million A673 cells stably transduced with dCas9-KRAB with or without the gRNA mix and were injected subcutaneously into the flanks of NSG mice. A third group (non-transduced cells) was added as an additional control. Prior to inoculation, the cells were resuspended in 100 µL of ice-cold serum-free media and BD Matrigel Matrix High Concentration (BD Bioscience) at a 3:1 ratio. 12 mice were used per group. The widths and lengths of the tumors were measured every 2 days. Tumor volumes were calculated using the modified ellipsoid formula; $V = [(\text{width}^2) \times ½ \times \text{length}]$. Mice were culled (n = 3 per group, early time point) on day 15 post-cell implantation. The remaining mice were monitored daily until tumors were >1000 $mm^3$, then were humanely sacrificed. All resected tumors were either snap-frozen in liquid nitrogen for RNA extraction or fixed in 4% paraformaldehyde overnight at 4 °C, washed two times with PBS, and preserved in 70% ethanol for paraffin embedding.

*dCas9-KRAB delivery in vivo using polymeric nanoparticles*

EWS xenografts were developed and monitored as stated above with A673 cells. Once tumors reached average volumes of 100 $mm^3$, mice were randomly allocated into 3 different groups (12 mice per group): Vehicle Control (PBS), GBA-polymer encapsulating dCas9-KRAB with no gRNA and GBA-polymer encapsulating dCas9-KRAB RNPs gRNA mix targeting *EWSR1-FLI1*. The RNP formulations were prepared as described above for the in vitro experiments, with OptiMEM substituted with PBS. Each injection (100 µL per animal) contained a final concentration of 70 pmol dCas9-KRAB complexed with GBA-polymer at a mass ratio of 1.5: 1 polymer to dCas9-KRAB protein. Injections were given intratumorally every two days to eventually receive a total of four injections. 24 h post-last injection, 3 mice per group were culled (early time point) on day 15 post-cell implantation. The remaining mice were monitored every two days, then daily until tumors were >1000 $mm^3$, when humanely sacrificed and the tumors collected and stored as described above.

In a parallel experiment, and to control for the specificity of the gRNAs, an GFP gRNA mix was used (see Table S7). Once tumors reached average volumes of 100 $mm^3$, 20 mice were divided into 3 different groups (n = 8 for GBA-polymer/dCas9-KRAB RNP with GFP gRNA, n = 7 for GBA-polymer/dCas9-KRAB RNP with no GFP gRNA, and n = 5 for Vehicle control (PBS). For injections were given intratumorally at 7, 9, 11, and 13 post-inoculations of A673 cells. Tumor size was monitored every two days then daily until mice were culled and tumors harvested as described above at end point when tumor sizes were >1000 $mm^3$.

*dCas9-KRAB expression after delivery of the polymeric nanoparticles in vivo*

For detection of dCas9-KRAB in mice tissues following RNP delivery, 36 NSG mice were divided into 3 groups (n = 12 per group). The mice were injected with a single intratumoral injection of either GBA-polymer/dCas9-KRAB RNP, GBA-polymer/dCas9-KRAB, or PBS. Animals (3 mice per group, per time point) were euthanized at time points 4, 8, 12 and 24 h post-intratumoral injection. Tissues were collected, washed in PBS, and fixed in 4% paraformaldehyde for histologic examinations.

*Propagation of EWS patient-derived xenograft (PDX) models*

The propagation of EWS PDX models were approved by the Animal Ethics Committee of the University of Western Australia (AE253 and AE256) and conducted in accordance with the Code of Practice for the Care and Use of Animals. EWS PDX tumor tissues were kindly provided by Professor Peter Houghton (Greehey Children's Cancer Research Institute, The University of Texas Health Science Center, San Antonio, USA). The EWS ES-4 PDX was subcutaneously transplanted as tumor tissue fragments (20–35 $mm^3$) in the right flank of 5-week-old female NSG mice. Mice weights and tumor volumes were monitored twice weekly for six months. Mice were excluded from the experiment if they did not demonstrate palpable tumors after six months, and the patient's tumors were considered as lacking engraftment.

36 mice were used to develop the first PDX passage (n = 6 mice). When tumor size reached nearly 800 $mm^3$, mice were humanly sacrificed, and tumors were harvested under sterile conditions. Resected tumors were utilized either to prepare FFPE blocks, make cryostat sections, or re-transplant into a new group of NSG mice to develop the second PDX generation (n= 5 mice). For making cryostat sections, two tumor pieces (2.5 × 2.5 × 5 mm) were preserved in sterile cryo-vials (ThermoFisher Scientific) in 1.5 mL freezing medium CryoStor® CS10 (Sigma-Aldrich) and stored in liquid nitrogen. To establish the third PDX passage, resected tumors were cut and placed into medium 199 (Gibco) on ice and reimplanted into 40 NSG mice per patient. When tumor volume reached 100 $mm^3$, mice were randomized into two groups (n = 10 mice/group) and animals were injected intratumorally with either GBA-polymer/dCas9-KRAB RNP, or GBA-polymer/dCas9-KRAB no gRNA. Mice received a total of 4 injections, once every two days. Tumor progression was monitored every two days using a Vernier caliper. Three mice per group were euthanized 4 h and 8 h after the last injection (early

time point), while the remaining mice were monitored every two days and then daily until end point. Tumors were collected and fixed for 24 h in 4 % paraformaldehyde for histological analysis or snap frozen and stored at -80°C for RNA extraction.

*Immunofluorescence staining of tissue sections*

Fixed tumors, cut sagittally into 4 μm sections and mounted onto slides, were deparaffinized with 3 x 5 min washes of xylene and rehydrated with 2 x 10 min washes of 100% ethanol and 95% ethanol. Antigen retrieval was performed using heated citrate buffer (10 mM sodium citrate, pH 6) for 10 min, permeabilized in permeabilization buffer (0.2% Triton X-100 in TBST) for 10 min, followed with a final wash of 2 x 5 min with TBST. The samples were blocked for 90 min at RT using blocking buffer (10% normal goat serum, 0.1% Triton X-100 in PBS). The slides were incubated with primary antibodies overnight at 4°C. The next day, slides were washed three times with PBS and then incubated with secondary antibodies Alexa Fluor 488 goat anti-mouse and Alexa Fluor 594 goat anti-rabbit secondary antibody (ThermoFisher Scientific) and Hoechst (Sigma-Aldrich) for 1 h. After mounting slides with SlowFade Diamond Antifade Mountant (ThermoFisher), images were captured using a fluorescent microscope (Nikon) and processed using NIS-Elements Software.

*Fluorescence in situ hybridization*

FISH was performed using EWSR1 Break Apart FISH Probe (Empire Genomics) following the manufacture's protocol. Briefly, deparaffinized tissue sections were incubated in 10mM citric acid pre-treatment buffer (pH 6.8) for 30 min at 90°C and then washed with 2x Saline-Sodium Citrate (2X SSC) wash solution for 5 min at RT. The slides were then placed in pepsin solution for 45 min at 37°C and then washed with 2X SSC for 5 min at RT. The tissue sections were dehydrated before the probes diluted in hybridization solution were applied. Slides were incubated for 7 min at 75°C for denaturation, followed by hybridization at 37°C in a humidified chamber for 24 h, and finally washed the following day using the wash buffer above. The tissue sections were then stained with DAPI and then mounted with SlowFade Diamond Antifade Mountant. Images were captured using a Nikon A1Si confocal microscopy and processed using NIS-Elements Software. For each tissue section, at least 100 nuclei were examined. Confirmed diagnosis of Ewing sarcoma was considered if >15% of cells had split signals.

*Statistical analysis*

Statistical analysis was performed using GraphPad Prism 8 software and are detailed in the figure legends. Unless otherwise specified, data were expressed as mean ± standard deviation (SD). A p-value of $p < 0.05$ was considered statistically significant. All experiments were carried out with three independent biological replicates. In RT-qPCR, mRNA abundance was normalized to no gRNA.
